# Design principles of collateral sensitivity-based dosing strategies

**DOI:** 10.1101/2021.03.31.437927

**Authors:** Linda B. S. Aulin, Apostolos Liakopoulos, Piet H. van der Graaf, Daniel E. Rozen, J. G. Coen van Hasselt

## Abstract

Collateral sensitivity (CS)-based antibiotic treatments, where increased resistance to one antibiotic leads to increased sensitivity to a second antibiotic, may have the potential to limit the emergence of antimicrobial resistance. However, it remains unclear how to best design CS-based treatment schedules. To address this problem, we use mathematical modelling to study the effects of pathogen- and drug-specific characteristics for different treatment designs on bacterial population dynamics and resistance evolution. We confirm that simultaneous and one-day cycling treatments could supress resistance in the presence of CS. We show that the efficacy of CS-based cycling therapies depends critically on the order of drug administration. Finally, we find that reciprocal CS is not essential to suppress resistance, a result that significantly broadens treatment options given the ubiquity of one-way CS in pathogens. Overall, our analyses identify key design principles of CS-based treatment strategies and provide guidance to develop treatment schedules to suppress resistance.

## Introduction

Antimicrobial resistance (AMR) is a worldwide health threat due to the reduction of clinically effective antibiotics. Current drug discovery pipelines of new-in-class antibiotic agents are insufficient to offset the emergence of new AMR^1^. Innovative strategies to reduce the rate that AMR develops are thus critically needed. Treatment with antibiotics in individual patients represents one important situation where *de novo* AMR may emerge^2,3^. However, current clinical antibiotic treatment strategies, *i.e*., which types of antibiotics are included as well as timing and dosage, typically do not explicitly consider within-host emergence of AMR. Instead, current strategies used in clinical practise are primarily based on exposure targets that are associated with sufficient bacterial kill in preclinical studies, or with clinical outcomes in patient studies^4^. Thus, there is need for clinical dosing strategies specifically designed to suppress AMR emergence^5^.

Trade-offs associated with AMR are of increasing interest to design antibiotic dosing strategies that suppress the within-host emergence of AMR^6^. In this context, collateral sensitivity (CS), where resistance to one antibiotic leads to increased sensitivity to a second antibiotic, has been proposed as a potential strategy to suppress AMR ^7,8^. CS has been characterized *in vitro*, typically by evolving AMR strains and then quantifying correlated changes in the sensitivity to other antibiotics^9–12^. CS effects have been characterized for several clinically relevant pathogens, including *Escherichia coli*^9^, *Pseudomonas aeruginosa*^13^, *Enterococcus faecalis*^14^, *Streptococcus pneumoniae*^15^, and *Staphylococcus aureus*^16^. CS relationships between antibiotics can either be one directional, where decreased sensitivity to one antibiotic show CS to a second antibiotic but not the reverse, or reciprocal, where decreased sensitivity to either of the antibiotics results in CS to the other. Reciprocal CS is often considered a prerequisite for effective CS-based treatments, but such relationships have been less frequently observed compared to one directional CS^9–16^.

CS-based treatment strategies can use different designs to combine antibiotics showing a CS-relationship, including simultaneous, sequential, or cyclic (alternating) administration. For example, consider a cycling drug strategy using two antibiotics showing reciprocal CS (Fig. 1). Initial treatment would start with antibiotic A. This leads to resistance to A and a corresponding increase in sensitivity to B. When treatment is switched to antibiotic B, the inverted selection pressure leads to the eradication of cells that are resistant to antibiotic A (due to CS), but possibly favouring any remaining cells that are resistant to B, but susceptible to antibiotic A. By cycling between the two drugs to sequentially eliminate all cells that show reciprocal CS, complete eradication can be achieved. Although the conceptual strategies of CS-based treatments have been discussed^6^, it remains unclear when CS-based dosing strategies are most likely to be beneficial, and how to design specific multi-drug antibiotic dosing schedules based on CS. Furthermore, it is unclear how pathogen-specific factors, such as CS effect magnitude and directionality, fitness costs of resistance, and mutation rates, as well as pharmacological factors related to pharmacokinetics and pharmacodynamics for different drug types, can affect optimal dosing schedules.

**Figure 1.**
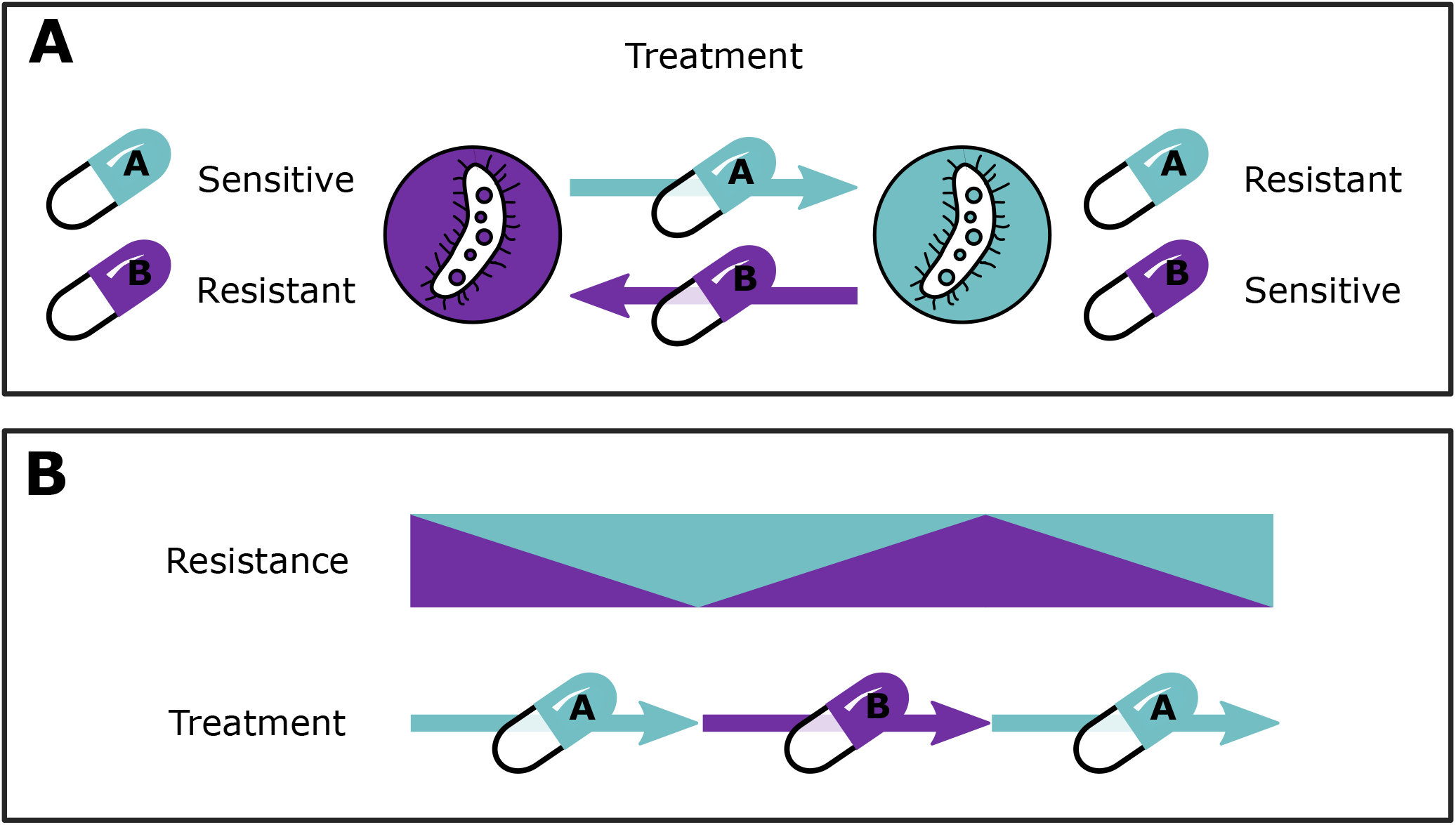
Concept figure of collateral sensitivity (CS)-based treatments using two hypothetical drugs, antibiotic A and B, based on Pál et al 2015 ^56^. **A**: Reciprocal CS relationship between antibiotic A and B. **B**: Theoretical cycling regimen exploiting CS between antibiotic A and B to supress resistance.

Experimental studies *in vitro* are essential to characterize the incidence, evolvability and magnitude of CS, all of which are important but isolated components that may contribute to the success of CS-based treatments ^9–16^. However, to translate *in vitro* CS findings to *in vivo* or clinical treatment scenarios, consideration of pharmacodynamic (PD) and pharmacokinetic (PK) factors is essential, as these determine the differential impact of different antibiotics on the concentration-dependent effects of bacterial growth, inhibition, and killing ^17,18^. By affecting bacterial dynamics, antibiotic PK-PD can have a profound influence on resistance evolution, and are therefore key elements to design optimised CS-informed treatments. In addition, it is necessary to disentangle the respective impacts of these separate parameters. Doing so requires a highly controlled system, where each factor can be modified separately; this this level of control cannot be established experimentally. To this end, a mathematical modelling approach can be highly valuable, as such models permit precise control of each factor. Additionally, mathematical models are important tools to integrate multiple biological and pharmacological factors contributing to treatment outcomes, including different PK parameters of specific antibiotics in patients, antibiotic-specific PD parameters, and pathogen specific characteristics such as strain fitness and the magnitude of CS effects. Thus, using a mathematical modelling approach allows us to address key questions relating to CS-based treatments that have yet to be fully answered.

A number of mathematical models have been developed to evaluate multi-drug therapies in relation to collateral effects often using shifts in MIC or other summary metrics as endpoints. These include deterministic^19^ and Markov^20^ models evaluating antibiotic cycling *in vitro* and *in silico*, which provide important insight into the importance of the design of the cycling regimen. Furthermore, a stochastic evolution model has been developed to assess the robustness of collateral sensitivity^21^. Despite the values of these models, they fail to characterise the bacterial dynamics underlying resistance evolution. Additionally, they do not include PK-PD relationships and lack consideration of clinical PK, which are key factors when translating the findings into clinical dosing strategies. Udekwu and Weiss developed a deterministic PK-PD model to study clinically relevant cycling schedules for CS-informed treatments and evaluated their ability to delay emergence of antibiotic resistance^22^. This simulation study serves as an important step toward designing clinically effective CS-based treatments. However, to take further steps towards such treatments, there is a need for a more comprehensive evaluation of the impact of several pharmacological and pathogen-associated factors related to dosing schedule designs, as well as specifically evaluate the impact of CS effects on treatment outcomes in comparison to the situation without CS.

In the current study we aim to build on previously established models in order to determine if and when CS-based dosing schedules lead to suppression of within-host emergence of antibiotic resistance. We utilise a mathematical modelling approach to comprehensively study the influence of key pathogen-specific factors and the contribution of PK and PD properties to identify key design principles to inform rational design of antibiotic multi-drug dosing schedules that suppress AMR.

## Methods

### Model framework

A differential-equation based model, consisting of components accounting for antibiotic PK and PD, and associated bacterial population dynamics, was developed to study the impact of differences in pathogen- and drug-specific characteristics for different treatment strategies using two antibiotics, hereafter referred to as drug A (D_A_) and drug B (D_B_). As a foundation for our model development, we used a deterministic PK-PD model developed by Udekwu and Weiss ^22^, which explores the impact of different multi-drug treatments on time to resistance development in the presence of CS. We advanced the model by incorporating mutations as random events to capture the stochasticity of resistance evolution. We integrated the different model components into a framework that allowed us to simulate antibiotic multi-drug treatments while altering drug- and pathogen-specific factors as a strategy to disentangle their impacts on resistance development.

#### Pharmacokinetics

A mono-exponential PK model was defined for both drugs D_*i*_, where *i* = {A,B}, as follows:

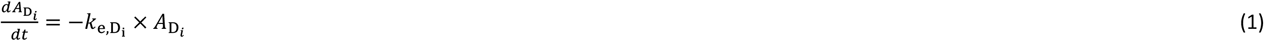

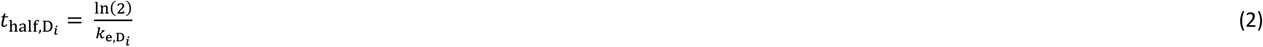

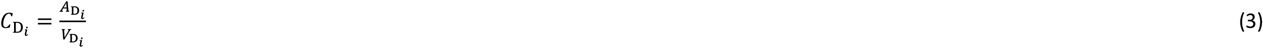

where Equation 1 describes the change of the amount of D_*i*_ in plasma over time after intravenous administration, 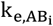 is the elimination rate of D_i_, which can also be expressed as a half-life 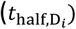 (Equation 2). The unbound plasma concentration 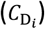, which is the assumed driver of the antibiotic effect, is calculated using the 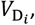, the distribution volume of D_*i*_ with the assumption of negligible protein binding (Equation 3).

#### Bacterial subpopulations

A model for antibiotic sensitive and resistant subpopulations was defined, comprising a four-state stochastic hybrid ordinary differential equation (ODE) model, where each state represents a bacterial subpopulation with different antibiotic susceptibility towards D_A_ and D_B_.

The model included an antibiotic sensitive bacterial subpopulation (WT) (Equation 4), one mutant subpopulation resistant to D_A_ but sensitive to D_B_ (R_A_) (Equation 5), one mutant subpopulation sensitive to D_A_ but resistant to D_B_ (R_B_) (Equation 6), and one double mutant subpopulation resistant to both D_A_ and D_B_ (R_AB_) (Equation 7). The initial bacterial population was assumed to be homogeneous and in the sensitive WT state unless stated otherwise.

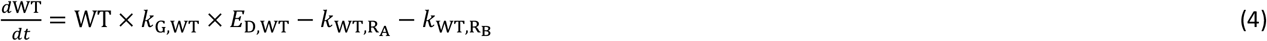

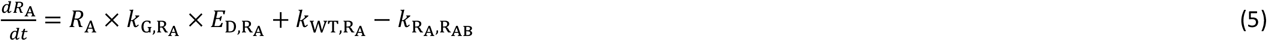

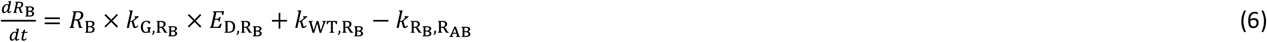

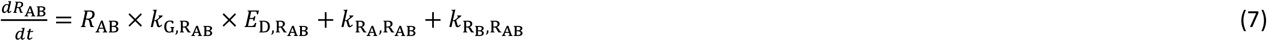

The above equations (Equation 4–7) describe the subpopulation specific rate of change for bacterial density, which is dependent on the bacterial density of subpopulation *z*, the subpopulation specific net growth (*k*_G,*z*_), the drug effect (*E*_D,*z*_), and mutation transition(s) (*k_z_*_,*M*_), if present.

#### Resistance mutation

Resistance evolution was included as stochastic mutation process. This process was modelled using a binomial distribution *B* with a mutation probability equal to the mutation rate (*μ*). The number of bacteria mutated per time step *k*_*z*,*M*_ depended on the number of bacteria available for mutation (*n*_z_), *i.e*., the bacterial subpopulation density of subpopulation z multiplied by the infection volume *V*, for mutation at time *t* (Equation 8). Double resistant mutants evolved through two mutation steps.

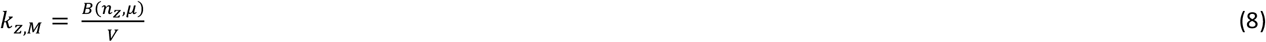

#### Pharmacodynamic effects

Drug effects on bacterial subpopulations (Equation 4–7) were assumed to be additive and the total drug effect for each subpopulation *z* (*E*_D,*z*_) was implemented as follows (Equation 9):

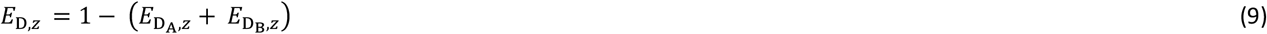

Here, antibiotic-mediated effects were implemented according to a PD model developed by Regoes *et al*. ^17^, where the effect of the *i*^th^ antibiotic on bacterial subpopulation 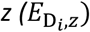 was related to the unbound drug concentration (*C_D,i_*) according to Equation 10.

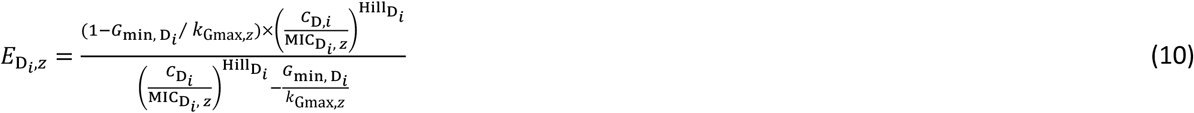

where 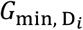 represents the maximal killing effect for the D_*i*_, *k*_Gmax,*z*_ is the subpopulation-specific maximal growth rate, 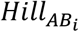 is the shape factor of the concentration-effect relationship, and 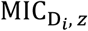 is the subpopulation-specific MIC of D_*i*_. This multi-parameter model allows for the description of the concentration-effect relationship of different shapes in relation to the subpopulation-specific MIC.

Sensitive bacteria were defined as having a MIC of 1 mg/L (MIC_WT_) and resistant as 10 mg/L (MIC_R_). Because the antibiotic concentrations are expressed as folds times MIC_WT_, the absolute value of MIC_WT_ is arbitrary. However, the ratio between MIC_WT_ and MIC_R_ is of relevance. A tenfold increase was chosen to represent a significant increase for a biologically plausible scenario. Resistance-related CS effects were included on the two single resistant mutants (R_A_ and R_B_), and were implemented as a proportional reduction (CS_A_ and CS_B_) of the MIC of the sensitive wild type bacteria (MIC_WT_). The double resistant mutant (R_AB_) was fully resistant (MIC = MIC_R_) to both antibiotic A and B, and did not have any collateral effects. The subpopulation- and antibiotic-specific MICs are stated below:

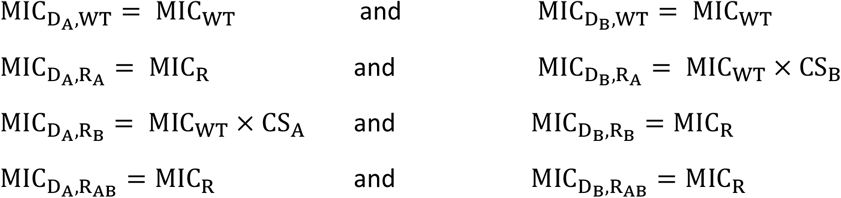

#### Growth rates and fitness effects

The maximal net growth rate (*k*_Gmax_) represents the maximal net growth of the wild type bacteria in the exponential growth phase. We considered resistance-associated fitness costs for the different mutant subpopulations. The fitness cost was incorporated using the factor *F*_fit_, which introduced a fractional reduction of *k*_Gmax_ for each resistance mutation. Thus, each subpopulation is associated with a specific maximal net growth rate (*k*_Gmax,*z*_), determined by the subpopulation-specific fitness, and was implemented according to Equation 11.

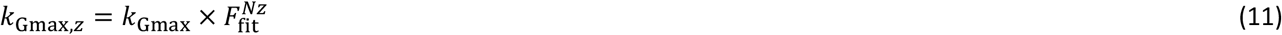

where 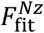 is the fitness cost factor per mutation, and *Nz* is the subpopulation-specific number of mutations (*Nz* = 0, 1 or 2).

The subpopulation-specific net growth in the absence of antibiotic (k_G,z_) was implemented according to Equation 12.

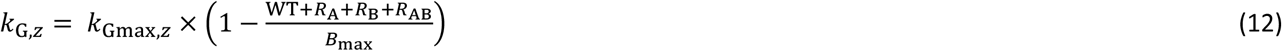

where *B*_max_ is the systems maximum carrying capacity, and WT, *R*_A_, *R*_B_, *R*_AB_ represent the bacterial densities of the four different subpopulations, respectively.

### Pathogen- and infection-specific parameters

The maximal growth rate (*k*_Gmax_) of the hypothetical pathogen was 0.7 h^−1^, thus representing a doubling time of one hour. The infection-specific parameters were chosen to represent a human bacteraemia, thus a typical human blood volume of five litres was used as the infection site volume^23^. An initial bacterial density of 10^4^ colony forming units (CFU)/mL was used to represent an early stage of an established infection^24^. A system carrying capacity limitation (*B*_max_) of 10^8^ CFU/mL^24^ was implemented according to Equation 12. When the maximum carrying capacity is reached, the net growth of the total bacterial population is zero, resulting in stationary phase. During this phase bacterial replication continues but is offset by bacterial death at the same rate, thereby still allowing for resistance mutations to occur. Resistance mutation rates of 10^−6^ and 10^−9^ mutations/base pair/hour were chosen to represent a high and a moderate mutation rate scenario, respectively^25^.

### Drug-specific parameters

The two hypothetical antibiotics used for the simulations (D_A_ and D_B_) have identical one-compartmental PK with distribution volumes of one litre, five-hour half-lives, and no protein binding. Their half-lives were selected to represent antibiotics with clinically relevant short half-lives, thereby rapidly reaching steady-state concentrations with minimal accumulation. The drugs were administrated as intravenous bolus doses twice daily over a treatment duration of two weeks. Several different dosing regimens were simulated including monotherapy, sequential non-repetitive dosing, one- and three-days repeated cycling regimens, and simultaneous dosing. Here, sequential non-repetitive dosing represents a multi-drug treatment using D_A_ for the first seven days and then switching to D_B_ for the remaining seven days of the treatment. The repeated cycling regimens represent multi-drug treatments starting with D_A_ for the duration of the cycling interval (*i.e*., one or three days), then switching to D_B_ for the same duration, and then back to D_A_, continuing the repeated cycling until the end of treatment. For sequential and repeated cycling treatments D_A_ was always used as the starting drug. The doses used were obtained by calculating the required dose to achieve appropriate average steady state concentration (C_ss_) relative to the MIC_WT_. The lowest dose (using 0.5 x MIC increments) that gave kill or stasis of the WT bacteria within the 24 hours of treatment, but allowed for resistance development during monotherapy, was selected for all dosing regimens except for the simultaneous dosing, for which the dose for the individual antibiotics were reduced by half in order to allow for resistance development. Four different PD types were included using different combinations of representative parameter values of Hill (driver of antibiotic effect) and *G*_min_ (type of antibiotic effect). The driver of the antibiotic effect, which is reflected by the steepness of the concentration-effect relationship (Hill), where shallow relationships are associated with time-over-MIC-dependency (Hill = 0.5) and steep relationship with concentration-dependency (Hill = 3). The type of antibiotic effects are commonly divided into bacteriostatic (*G*_min_ = −1) or bactericidal (*G*_min_ = −3). The corresponding PK-PD relationship of the four different antibiotic types is shown in Fig. 3.

**Figure 2.**
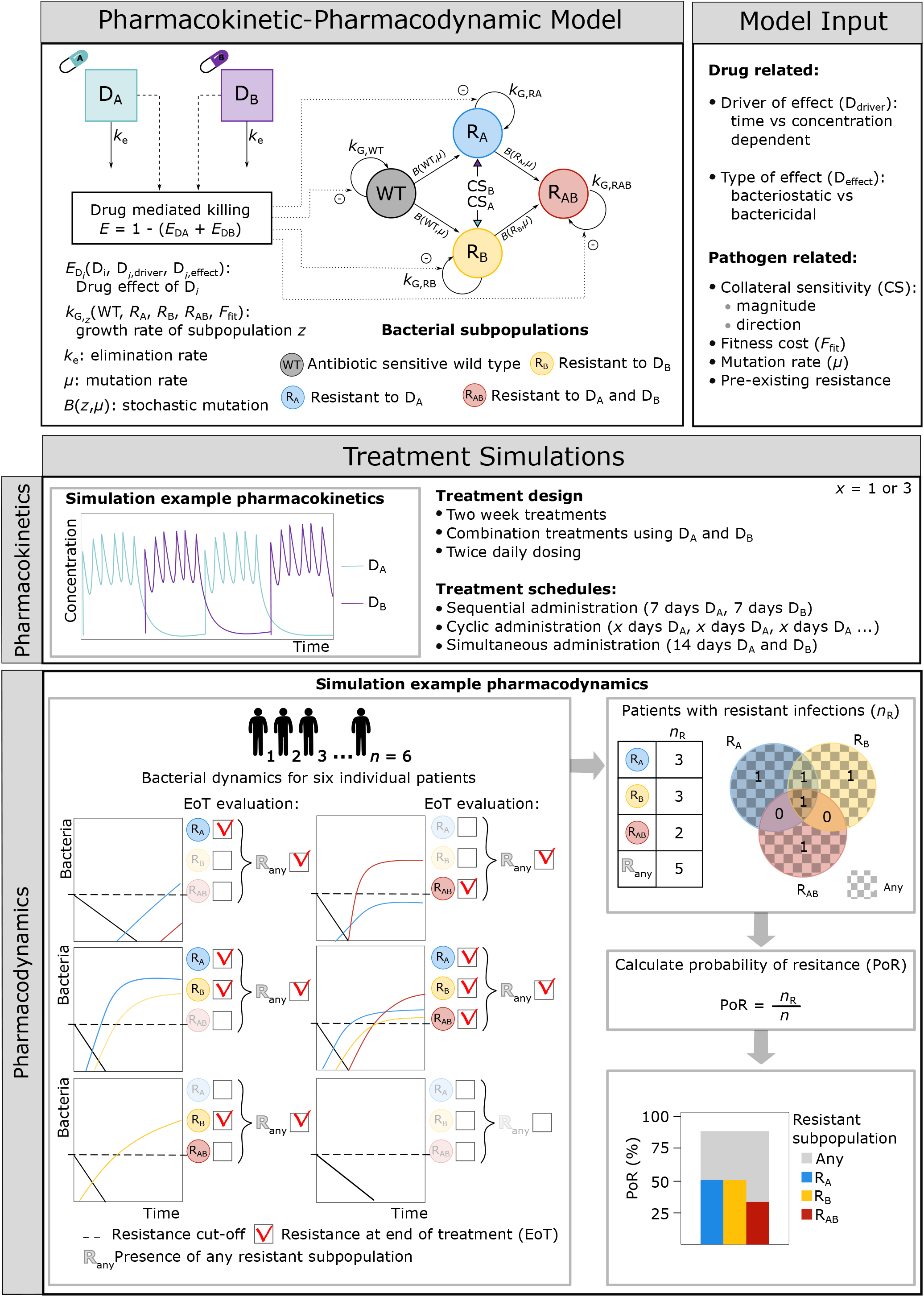
Simulation workflow. Pharmacokinetic-pharmacodynamic (PK-PD) framework comprised of four bacterial subpopulations (WT, R_A_, R_B_, R_AB_) and the PK-PD of two hypothetical drugs (D_A_ and D_B_). The framework includes fixed infection- and pathogen-specific parameters and fixed drug PK parameters. The model input includes both drug- and pathogen-related factors, which vary between different scenarios. The framework was used to simulate different treatment schedules of two-week multi-drug treatments using D_A_ and D_B_ for n patients. In the example a three-day cycling treatment regimen (PK panel) is simulated for six patients. The resulting patient-specific bacterial profiles are shown in the PD panel. Resistance was evaluated for each patient and bacterial subpopulation at the end of treatment (EoT), for which the corresponding probability of resistance (PoR) was calculated.

**Figure 3.**
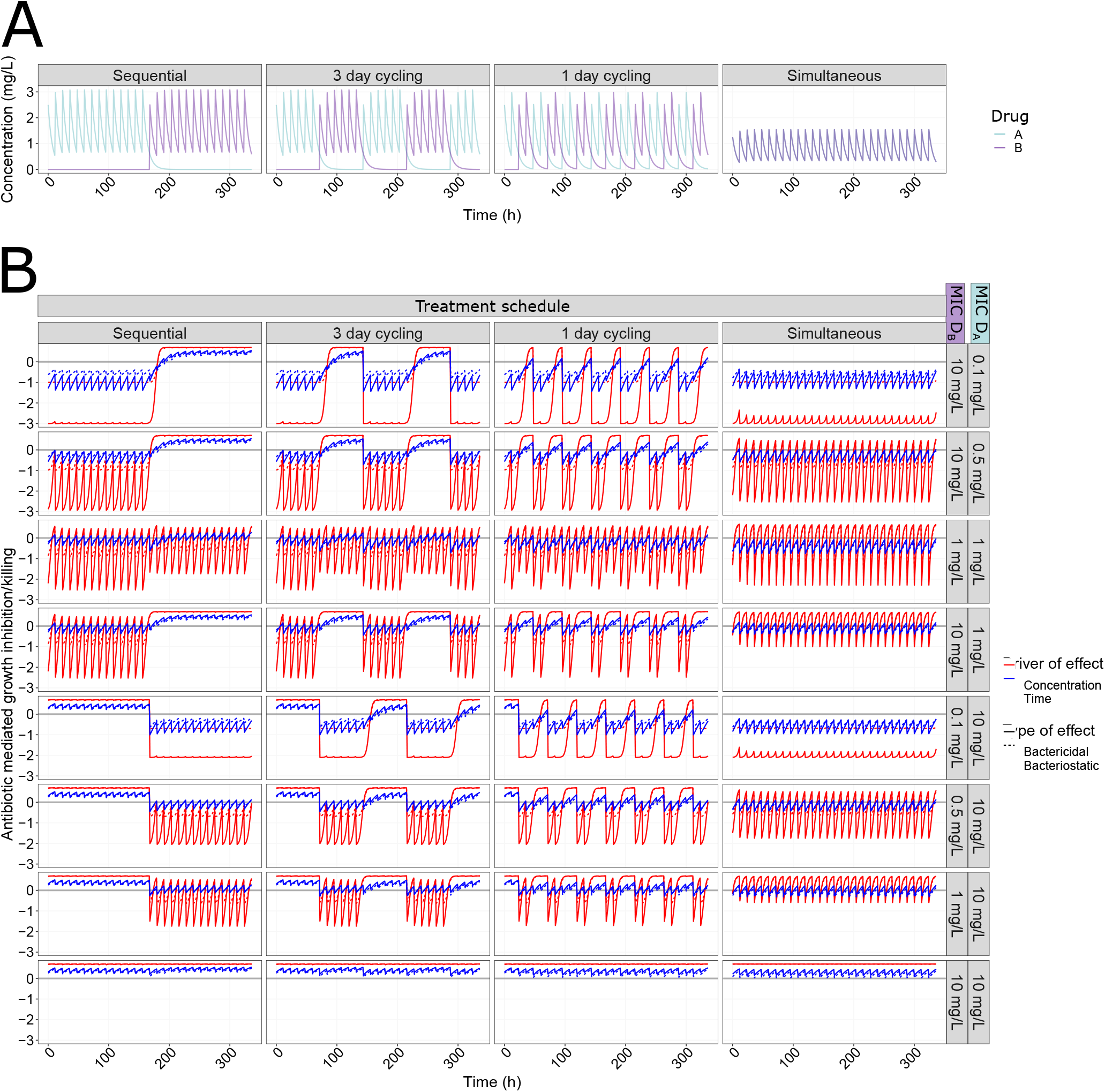
MIC-specific PK-PD relationships. **A**: Initial pharmacokinetic (PK) profiles of mono or multi-drug treatments using two hypothetical antibiotics D_A_ (turquoise) and D_B_ (purple) where both drugs follow one-compartmental kinetics with first order elimination. The drugs were administrated intravenously twice daily according to four different treatment schedules (columns), including non-reparative sequential administration, repetitive cycling administration, and simulations administration. Dosages used were related to average steady state concentration of 1.5 mg/L (1.5 x MIC_WT_) or 0.75mg/L (0.75 x MIC_WT_) for simultaneous dosing. **B**: Pharmacodynamic profiles related to different treatment schedules using different antibiotic drug types including concentration-(Hill = 3, red) or time-(Hill = 0.5, blue) dependent antibiotics and bactericidal (*G*_min_ = −3, solid) and bacteriostatic (*G*_min_ = −1, dashed), where the effect is representing the proportional bacterial growth inhibition/killing of different bacterial phenotypes (rows). The bacterial phenotypes are associated with different sensitivities towards D_A_ and D_B_. The effect is driven by the PK profile shown in panel A according to Equation 9 and 10.

### Simulation scenarios

An initial set of dose-finding simulations revealed that monotherapy required C_ss_ equal to 1.5 mg/L (1.5 x MIC_WT_) to achieve killing of the WT while allowing for emergence of resistance in the absence of CS, regardless of the drug type used (Supplementary Fig. 1). These dosing conditions allow us to evaluate the effect of CS for the majority of the simulated treatments. However, for the treatments where antibiotics were dosed simultaneously, half of the dose (C_ss_ 0.75 mg/L) was used in order to keep the total dose constant, and to allow for resistance emergence in the absence of CS and comparable to non-simultaneous dosing regimens.

We used a systematic simulation strategy to study the impact of CS, fitness cost, mutation rate, and initial subpopulation heterogeneity in antibiotic sensitivity on the probability of resistance (PoR) development for different treatments. An overview of all simulated scenarios can be found in Supplementary Table 1. We simulated treatments using two same-type antibiotics (*G*_min,A_ = *G*_min,B_ and Hill_A_ = Hill_B_) for scenarios without CS as well as in the presence of one directional and reciprocal CS in the magnitude of 50% or 90% (2 or 10-fold) reduction of the sensitive MIC (Supplementary Table 1, Scenario 1 and 2). For these scenarios the resistance was assumed to occur without any fitness cost, thus allowing us to evaluated CS-specific effects on PoR. We also simulated a set of treatment scenarios using two different antibiotic types in the presence or absence of CS (Supplementary Table 1, Scenario 3). To assess the impact of therapeutic window of antibiotics, as reflected by the fold-difference of steady state concentration (C_ss_) to the MIC_WT_, we simulated different dosing levels resulting in a range of C_ss_ of 0.5-5 x MIC_WT_ (Supplementary Table 1, Scenario 4). Additionally, we simulated same-type treatment scenarios covering a wide range of fitness costs (10% to 50% fitness cost per mutation compared to the wild type) implemented as a growth rate reduction (Supplementary Table 1, Scenario 5). In order to better understand the interplay between CS and fitness cost we simulated these scenarios with and without CS. We further investigated how low levels of pre-existing resistance (1%) towards either AB_A_ or AB_B_ affected the PoR at the end of treatment for different dosing regimens (Supplementary Table 1, Scenario 6). Finally, we examined the effect of increased mutation rates on resistance development (Supplementary Table 1, Scenario 7).

Each simulated scenario was realized 500 times (n), thus representing 500 virtual patients for which the within-patient resistance development was assessed. For each scenario we evaluated different multi-drug treatments regimens, including within-patient cycling and simultaneous administration. We note that most previously conducted studies investigating the clinical utility of antibiotic cycling and mixing to supress AMR have evaluated stewardship strategies at a community level^26–28^, *e.g*., between patients within a hospital ward. However, community level strategies are conceptually different from the within-patient multi-drug treatment strategies we investigate in this analysis. Therefore, the results we derive from our simulations are not directly comparable to the findings from such epidemiological studies.

### Evaluation metrics

We computed the probability of resistance (PoR), which was defined as resistant bacteria reaching, or exceeding, the initial bacterial density of 10^4^ CFU/mL at the end of treatment, for each subpopulation separately (Equation 13)

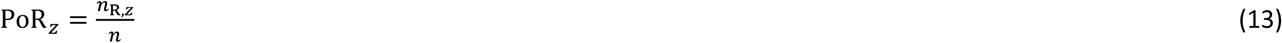

where *n_R,z_* denotes the number of patients having resistant bacteria of subpopulation *z* at the end of treatment (Equation 14):

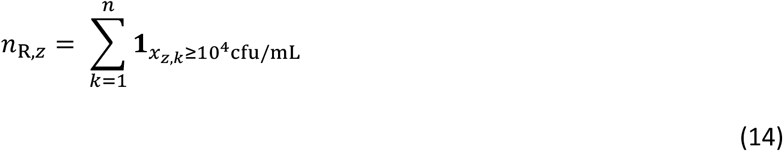

where **1** denotes the indicator function and *x_z,k_* denotes the bacterial density of subpopulation *z* at the end of treatment of patient *k*.

We also calculated the PoR for the case where any, *i.e*., one or more, resistant subpopulation(s) exceeded the resistance cut-off (R_Any_).

The standard error (SE) of the PoR was calculated according to Equation 15.

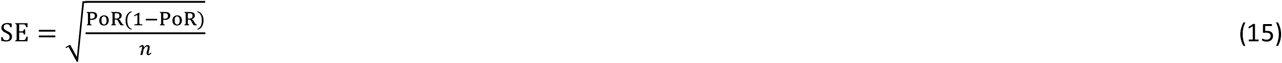

### Software and model code

All analyses were conducted in R version 4.0.5, using the ODE solver package RxODE (version 1.0.0-0)^29,30^. The associated model code is available at https://github.com/vanhasseltlab/CS-PKPD)^31^.

## Results

### Drug type and treatment schedule influence the probability of resistance

We simulated multi-drug antibiotic treatments using two antibiotics of the same type, with either no (0%) or reciprocal CS (50 or 90% decrease comparing to MIC_WT_). We show that the impact of reciprocal CS on resistance dynamics is dependent on the simulated drug type and dosing regimen (Fig. 4). In our simulations, treatments with concentration-dependent antibiotics could achieve full CS-based resistance suppression for dosing schedules using one-day cycling interval (Fig. 4C, 4G) or simultaneous administration (Fig. 4D, 4H). A 50% MIC reduction was sufficient to achieve this effect for all of the four treatments, which is relevant in light of experimental results consistent with these CS magnitudes^15,16,20,32,33^. Treatments using time-dependent antibiotics dosed according to these schedules (Fig. 4K, 4L, 4O, 4P) were efficient in fully supressing resistance with or without CS. Full resistance suppression was not achieved by any of the other treatment schedules tested. Although none of the CS-based treatments dosed according to the three-day cycling regimen managed to fully supress resistance, the ones using time-dependent antibiotics (Fig. 4J, 4N) did show reduced PoR in the presence of CS. For these treatments, the effect of CS was most prominent for bacteriostatic antibiotics (Fig. 4N) where a CS magnitude of 90% resulted in a decrease of the PoR of12.6% for R_Any_. Importantly, we also find that for some treatments the presence of CS was not only unable to fully supress resistance, but favoured the formation of double resistant mutants (Fig. 4F, 4I, 4M).

**Figure 4.**
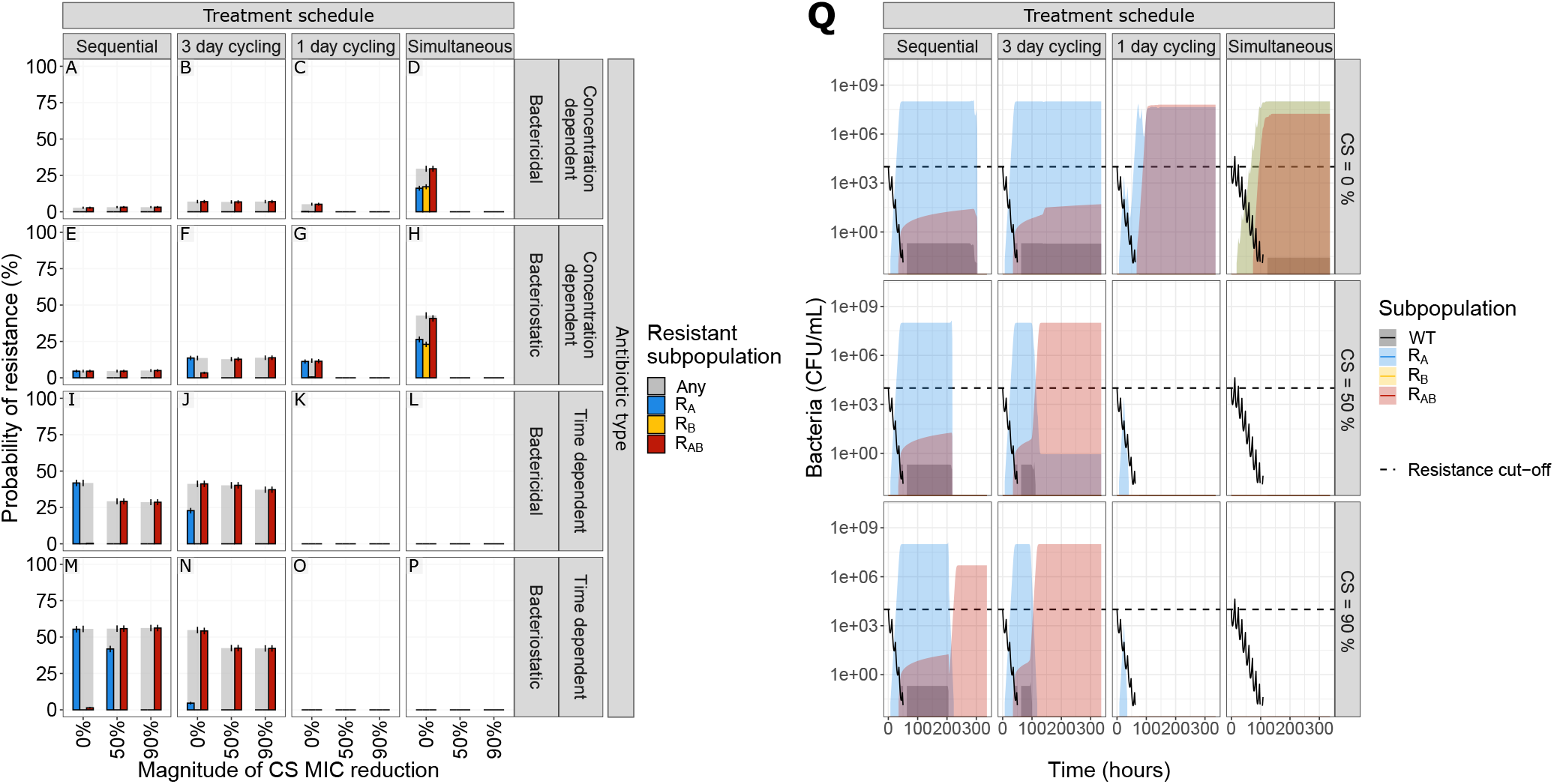
The effect of treatment design and antibiotic in relation to different levels of collateral sensitivity (CS) on the probability of resistance (PoR) at end of treatment. The simulations show that CS had a profound impact on resistance development for treatments with concentration dependent antibiotics with one-day cycling interval or simultaneous administration. **A-P**: PoR was estimated at end of treatment for treatments using different designs (columns) and antibiotic types (rows), where each simulated scenario was realized 500 times. Subpopulation-specific PoR are indicated by different colour and R_Any_, defined as the presence of any resistant subpopulation, is indicated in grey. Data are presented as mean PoR with the error bars represent the standard error of the estimation. **Q**: Bacterial dynamics relating to different treatment schedules using concentration-dependent bacteriostatic drugs, where each simulation was realized n=500 times. Subpopulation-specific bacterial density are indicated by different colours, where the solid lines indicate the median and the shaded area covers the 5^th^-95^th^ percentiles of the predictions. The resistance cut-off (dashed line) is used for end of treatment evaluation of resistance.

### Directionality of CS effects influence the probability of resistance

We next sought to determine if reciprocity is a requirement for CS-based treatments to suppress *de novo* resistance. We find that bactericidal and bacteriostatic drugs showed the same overall behaviour for treatment outcomes when tested in relation to CS directionality (Supplementary Fig.2). We specifically focus on the one-day cycling and simultaneous treatment that appeared to be most successful in fully supressing resistance for reciprocal CS. We find that for the one-day cycling regimen the presence of one directional CS for the second administrated antibiotic (D_B_) is sufficient to fully suppress resistance development. This is illustrated for treatments using concentration-dependent bacteriostatic antibiotic in Fig. 5. In this scenario, one directional CS results in resistance levels close to the reciprocal scenario (*e.g*., one directional 50% CS resulted in 0.4% PoR of R_Any_ for bacteriostatic (Fig. 5A) vs 0% for reciprocal CS (Fig.5B)). In contrast, when CS is only present for the first antibiotic administered (D_A_), we found resistance levels close to the scenario without any CS (PoR 11.5% (Fig. 5D) vs 12.4% (Fig. 5C)). Overall, these results suggest that when using a drug-pair without reciprocity, the order of administration has a large impact on treatment success and that therapy should be initiated with the antibiotic for which there is no CS. This strategy allows for evolution and growth of R_A_ on the first day, while R_B_ is supressed by D_A_. When the selection is inverted on day two, the low levels of R_A_ are effectively killed by D_B_ in the presence of CS. In the absence of CS towards D_B_, R_A_ will reach high levels, which can lead to further evolution of R_AB_. When simultaneous administration of concentration-dependent antibiotics is used, we found that reciprocity is necessary to fully supress resistance, as one directional CS will only supress resistance for the resistant subpopulation which shows CS (Fig. 5A and 5B). However, one directional CS did reduce the PoR for R_Any_ by approximately 50% (ΔPoR −19.6% and −19.2% for CS_A_ and CS_B_, respectively) for both of these treatments.

**Figure 5.**
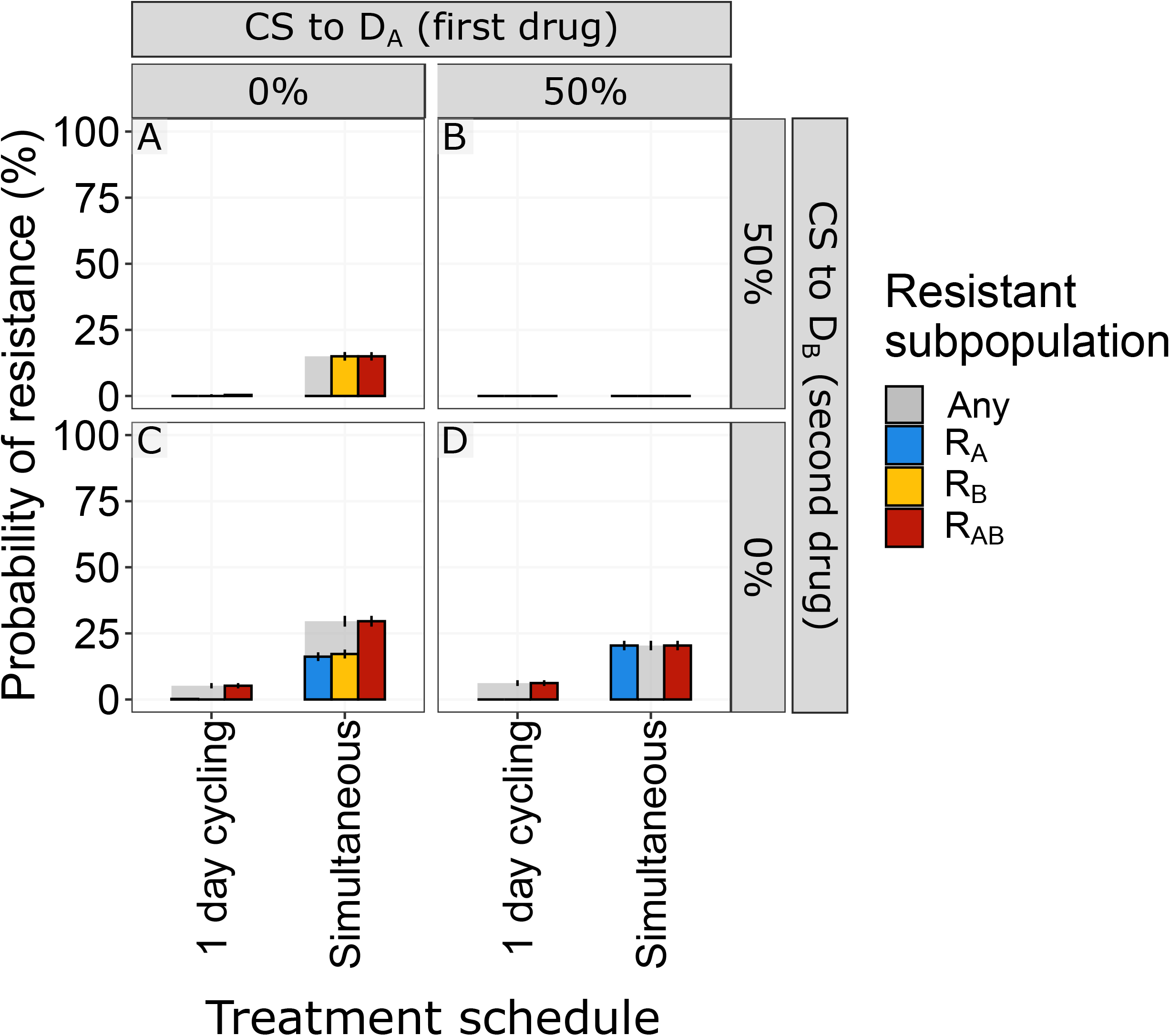
The effect of the direction or reciprocally of collateral sensitivity (CS) on end of treatment probability of resistance (PoR). PoR was estimated at end of treatment for different CS scenarios using concentration dependent bacteriostatic drugs. Subpopulation-specific PoR is indicated by different colour and R_Any_ resistance, defined as the presence of any resistant subpopulation, is indicated in grey. Each simulated scenario was realized n=500 times. Data are presented as mean PoR with the error bars represent the standard error of the estimation. For the one-day cycling regimen it became evident that the CS towards the second administrated drug (D_B_) was driving the effect, CS-based dosing using simulations administration of concentration dependent antibiotics showed that reciprocity is necessary to supress overall resistance.

### Administration sequence and antibiotic type influence resistance suppression

As CS does not only occur between antibiotics of the same type, it is important to understand how the administration sequence of different-type antibiotics affects resistance evolution. Our results for one-day cycling and simultaneous schedules demonstrated that the suppression of *de novo* resistance was mainly driven by the first administered antibiotic (D_A_) for all non-simultaneous regimens (Supplementary Fig. 3), highlighting the importance of drug sequence. In line with our findings for multi-drug treatments using same-type antibiotics (Fig. 4), resistance was fully suppressed from CS only when using one-day cycling or simultaneous administration dosing regimens. Particularly for one-day cycling regimens (Fig. 6), initiating treatment with a time-dependent antibiotic was more effective at supressing resistance in the presence of reciprocal CS compared to the initial administration of a concentration-dependent antibiotic.

**Figure 6.**
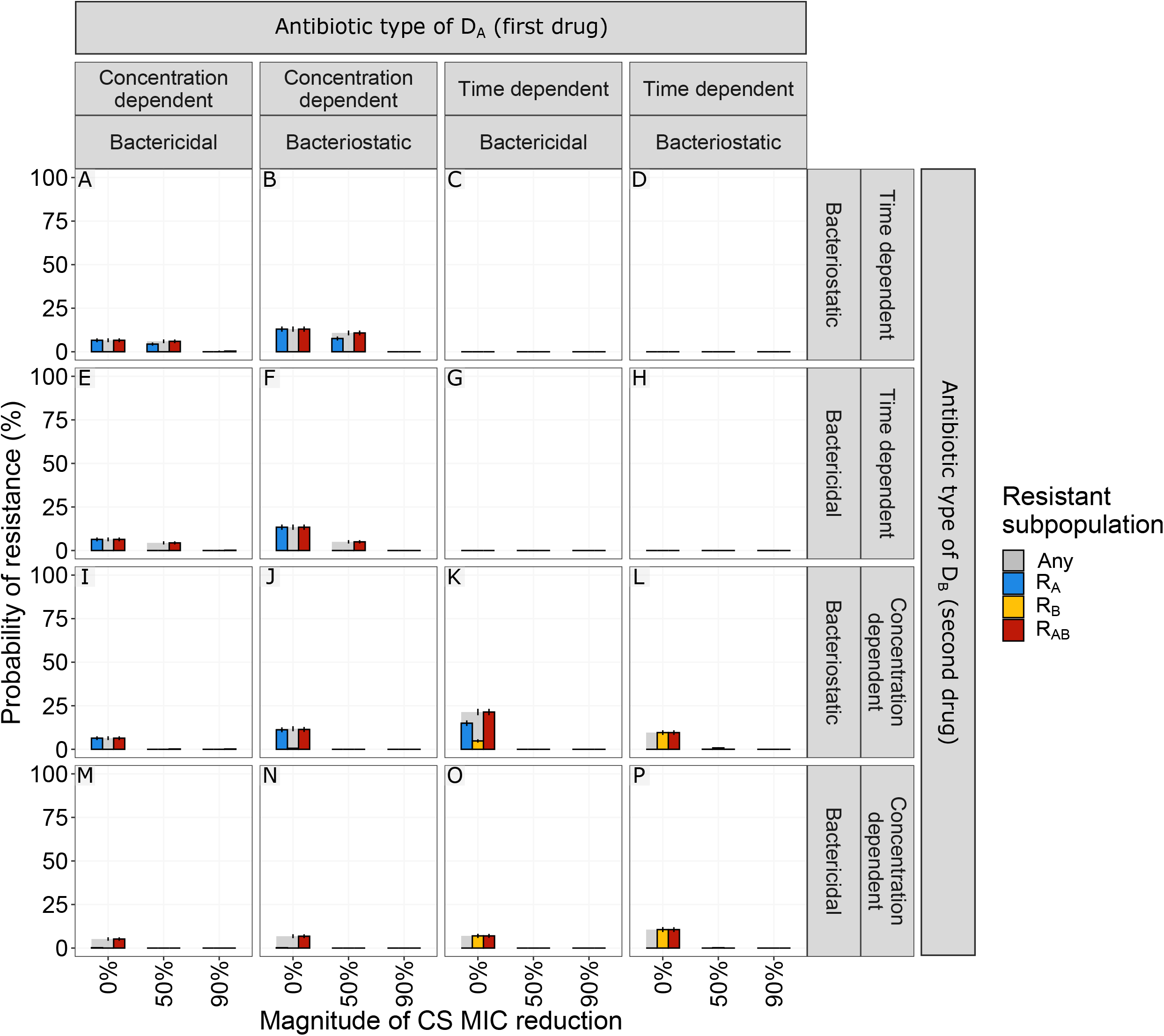
The effect of using different antibiotic types during one-day cycling multi-drug treatments in relation to different levels of collateral sensitivity (CS) on the probability of resistance (PoR) at the end of treatment. The simulations showed that initiating treatment with a time-dependent antibiotic was more effective in supressing resistance than with a concentration-dependent antibiotic in the presence of reciprocal CS. Each simulated scenario was realized n=500 times. PoR was estimated at the end of treatment for one-day cycling regimen with different antibiotic combinations. Subpopulation-specific PoR is indicated by different colour and R_Any_, defined as the presence of any resistant subpopulation, is indicated in grey. Data are presented as mean PoR with the error bars represent the standard error of the estimation.

### CS-based multi-drug treatments show greatest promise for antibiotics with a narrow therapeutic window

Although many antibiotics are well-tolerated and can be dosed well above the MIC of susceptible strains others, *e.g*., aminoglycosides, display a narrow therapeutic window due to toxicity^34–36^. Understanding the relationship between average steady-state concentrations (C_ss_) and the impact of CS on *de novo* resistance development would help identify in which clinical scenarios CS could be exploited to improve treatment. To this end, we simulated a set of dosing regimens (using same-type antibiotics) resulting in C_ss_ ranging between 0.5-5 x MIC_WT_. These simulations revealed that CS has the greatest impact on R_Any_ for C_ss_ close to the MIC_WT_ (Fig. 7 and Supplementary Fig. 4). Most treatments showing a benefit of CS lost the advantage when the C_ss_ exceeded 1.5 x MIC_WT_. The only exception was one-day cycling treatment using concentration-dependent bacteriostatic drugs, which retained an advantage up to C_ss_ of 2 x MIC_WT_ (Fig. 7G).

**Figure 7.**
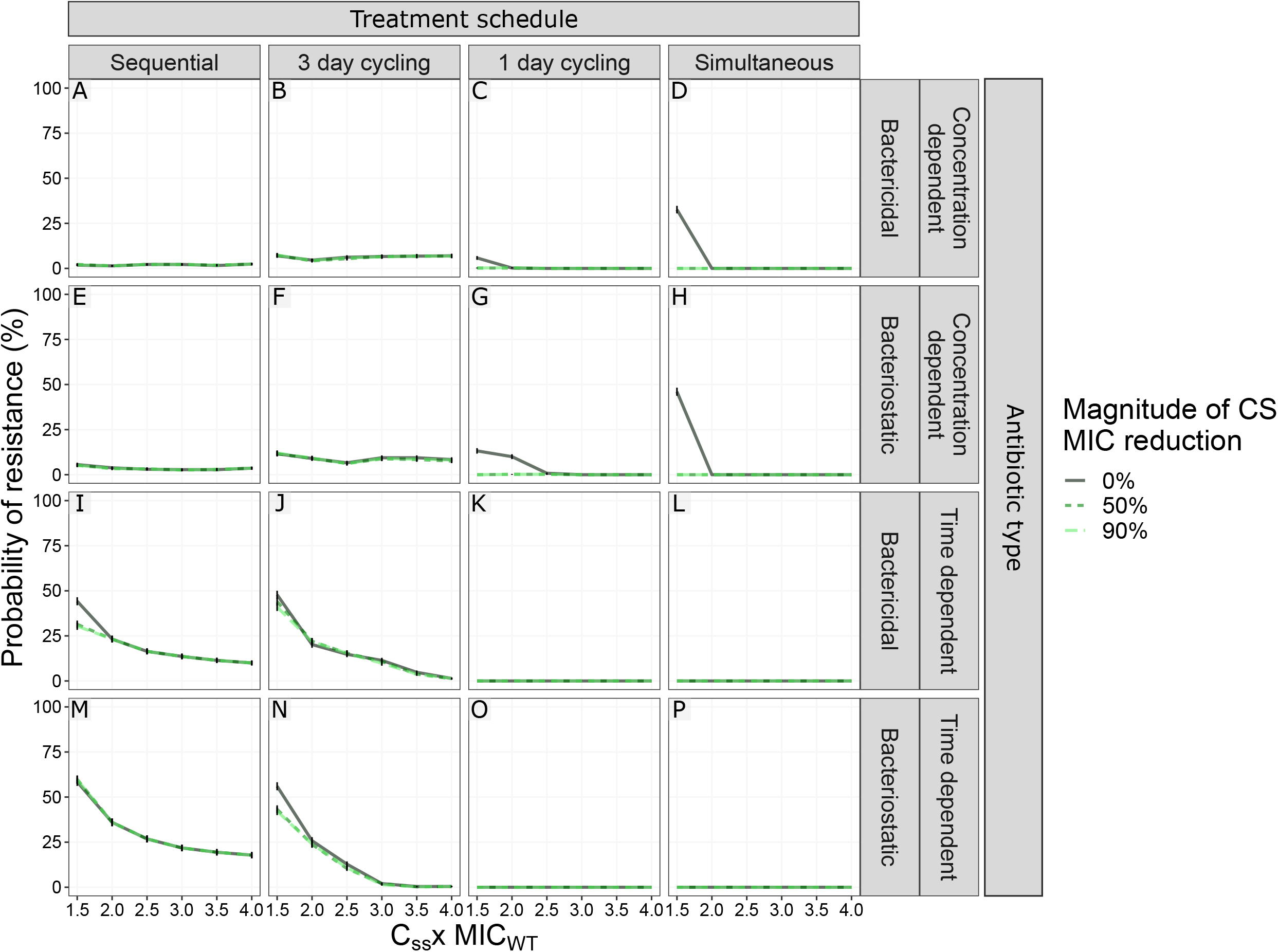
The effect of antibiotic steady state concentrations (C_ss_) in relation to different levels of collateral sensitivity (CS) on the probability of resistance at the end of treatment (PoR). The simulation revealed that CS had the largest impact on PoR for C_ss_ close to MIC of the wild type strain (MIC_WT_). C_ss_ was expressed as factor difference from the MIC_WT_. For the dosing regimen using simultaneous administrated antibiotics the C_ss_ represent the total antibiotic C_ss_, where the individual antibiotics were dosed at 0.5 x C_ss_. PoR of R_Any_, defined as the presence of any resistant subpopulation, was estimated at the end of treatment for treatments using different designs (columns) and antibiotic types (rows). Each simulated scenario was realized n=500 times. Colour and line-type indicate the magnitude of reciprocal CS simulated. Data are presented as mean PoR with the error bars represent the standard error of the estimation..

### Fitness cost of antibiotic resistance can contribute to the success of CS-based treatments

Resistance evolution is commonly associated with fitness costs^37^. We studied the impact of different levels of fitness cost on the suppression of *de novo* resistance development (Fig. 8). Fitness cost was included as a fractional reduction of growth per mutation, thereby doubly penalising the double resistant mutant R_AB_. In the absence of CS, fitness cost below 50% per mutation had little impact (|ΔPoR| ≥ 5%) on R_Any_ for most treatment scenarios. However, when concentration-dependent bactericidal drugs were dosed simultaneously the presence of fitness costs slightly increased the PoR (maximum ΔPoR 10.2 %). The presence of fitness cost increased the impact of CS on PoR the three-day cycling regimen using time dependent antibiotics (Fig. 8J and 8N), which failed to fully supress resistance in the presence of fitness cost-free CS. The fitness cost generating the largest impact of CS for these treatments on PoR was 40% and 50% cost per mutation when treated with bacteriostatic (Δ PoR −48.4) and bactericidal (Δ PoR −42.8%) drug, respectively.

**Figure 8.**
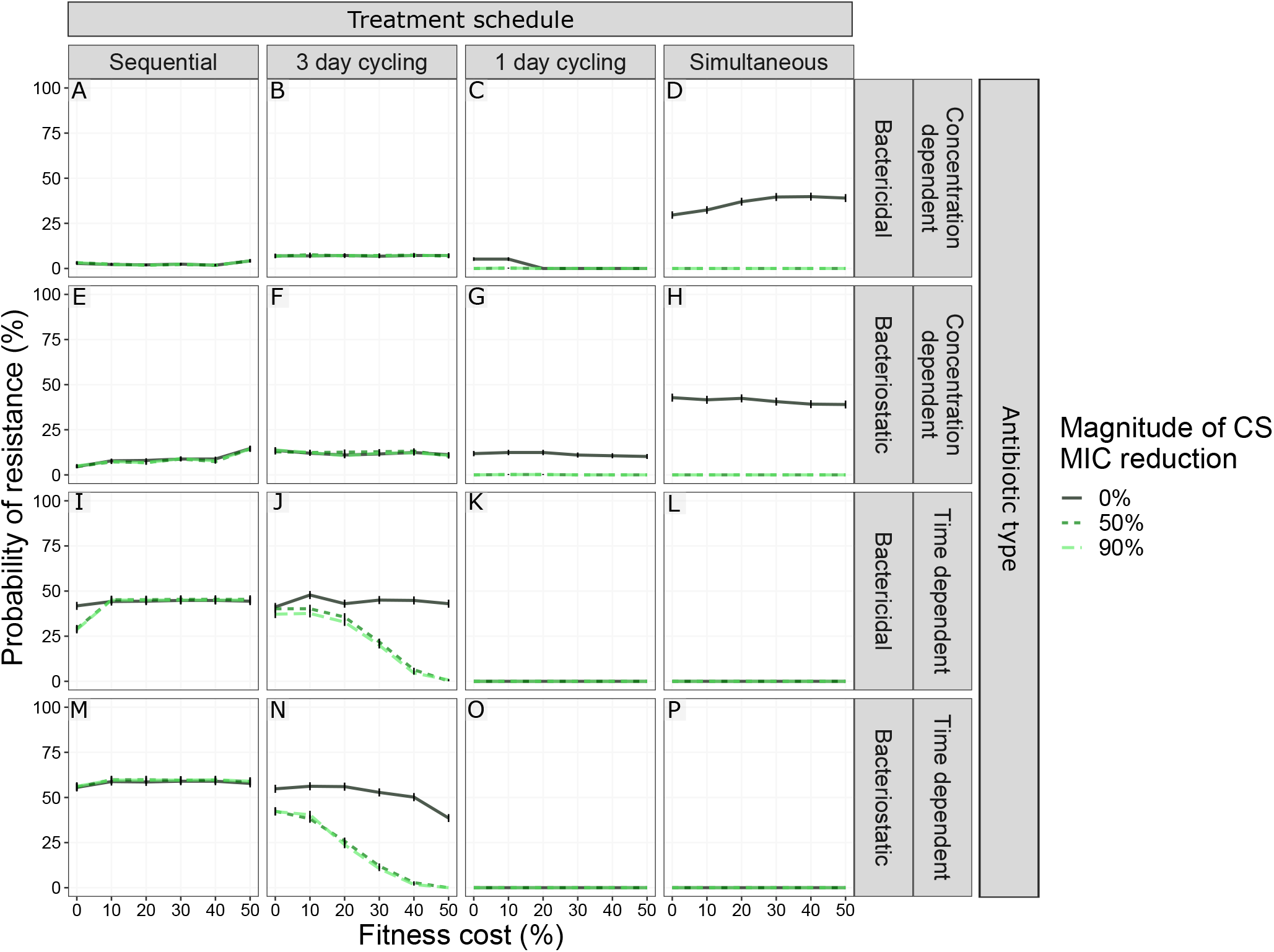
The effect of fitness costs for developing resistance for different levels of collateral sensitivity effects on the probability of resistance (PoR). PoR of R_Any_, defined as the presence of any resistant subpopulation, was estimated at end of treatment for treatments using different designs (columns) and antibiotic types (rows). Colour and line-type indicate the magnitude of reciprocal CS simulated. Each simulated scenario was realized n=500 times. Data are presented as mean PoR with the error bars represent the standard error of the estimation.

### CS-based simultaneous treatment designs suppress pre-existing resistance

The presence of rare pre-existing resistant cells in the bacterial population establishing an infection is clinically associated with antibiotic-treatment failure^38^. We here studied if CS-based dosing schedules can be used to eradicate such a heterogeneous population (Fig. 9 and Supplementary Fig. 5). In the absence of CS, most of the simulated treatment scenarios resulted in a higher probability of the expansion and fixation of pre-existing resistant sub-populations. As with *de novo* resistance and cycling regimens, the benefit of reciprocal CS was only apparent when resistance was towards the second antibiotic (subpopulation R_B_). This is illustrated with the one-day cycling treatments shown in Fig.9, where all CS-based treatments could supress PoR for pre-existing R_B_, but failed for all with pre-existing R_A_. For three-day cycling regimens and pre-existing resistance towards the first antibiotic, CS was shown to increase the PoR for R_AB_ (Supplementary Fig. 5). In the presence of CS, all simultaneously dosed treatments were effective in fully suppressing resistance regardless of pre-existing resistance (Supplementary Fig. 5).

**Figure 9.**
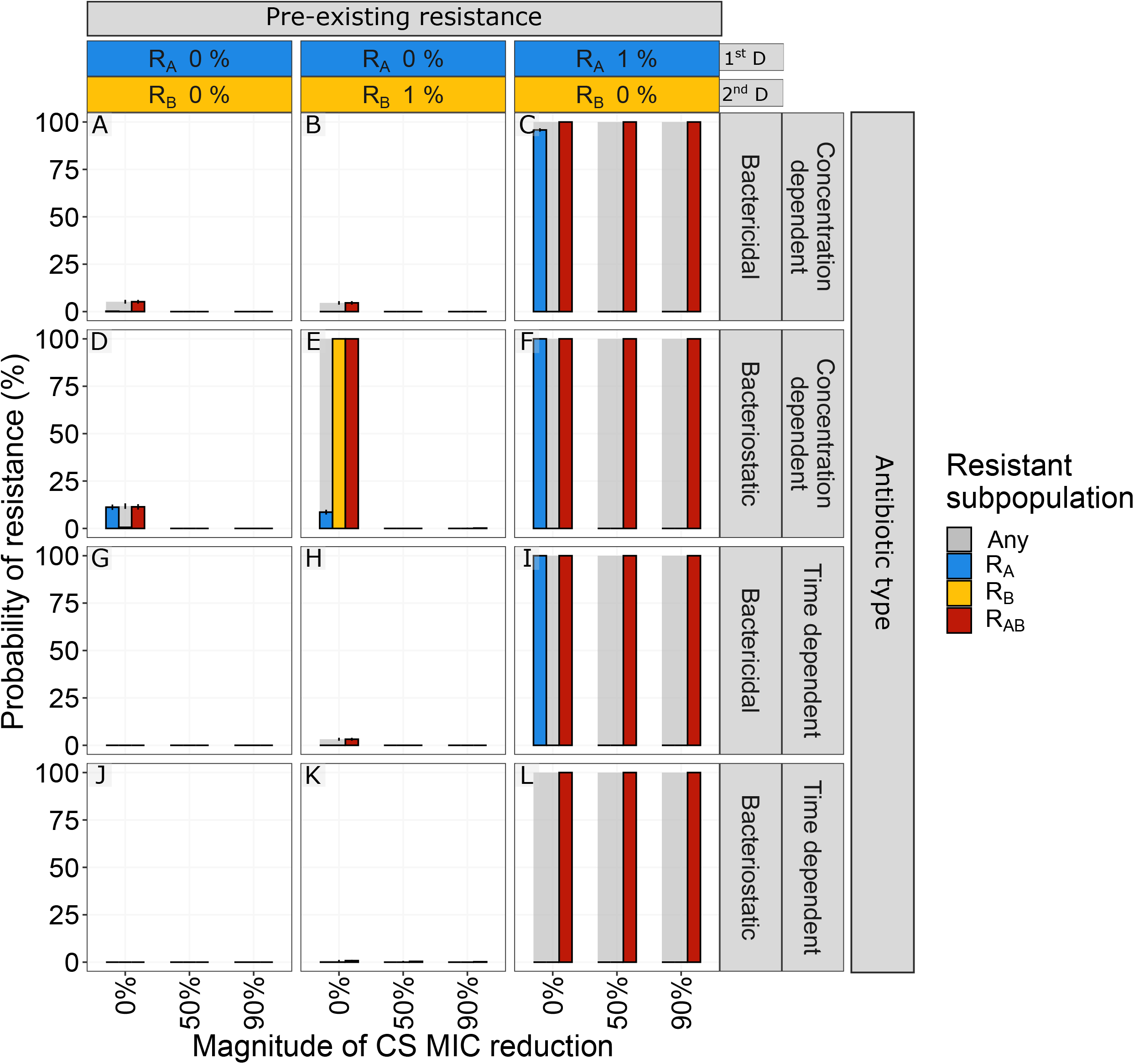
The effect of pre-existing resistant mutants for different magnitudes of collateral sensitivity on the probability of resistance (PoR). PoR was estimated at the end of treatment for different scenarios of low levels of pre-existing resistance (columns) and antibiotic types (rows). Subpopulation-specific probability of resistance is indicated by colour and PoR of R_Any_, defined as the presence of any resistant subpopulation, is indicated in grey. Each simulated scenario was realized n=500 times. Data are presented as mean PoR with the error bars represent the standard error of the estimation.

### The combined effect of CS and mutation rate on resistance development differs between treatments

Because some antibiotic treatments can enhance the genome-wide mutation rate in pathogenic bacteria ^39^, we included a set of simulations with higher mutation rates than 10^−9^ mutations/bp/h (10^−8^-10^−6^ mutations/bp/h). We show that the impact of mutation rate on the PoR was dependent on the combination of treatment design and the antibiotic type used, especially in the presence of CS (Fig. 10). The largest impact of the interaction between CS and mutation on PoR was found for the extremes of the antibiotic switching time, *i.e*., one-day cycling and sequential treatment design (maximum ΔPoR −57.8% and −52.4%, respectively). In the absence of CS, an increased mutation rate generally led to an increased PoR, with the exception of simultaneous administration of time-dependent antibiotics, which actually resulted in full suppression of resistance regardless of CS and mutation rate. For sequential treatments using time-dependent antibiotics with reciprocal CS (Fig.10I, 10M), the highest PoR was observed at a mutation rate of 10^−7^ mutations/bp/hour, and decreased at higher mutation rates. For all mutation rates and in the presence of CS, simultaneous treatments supressed resistance.

**Figure 10.**
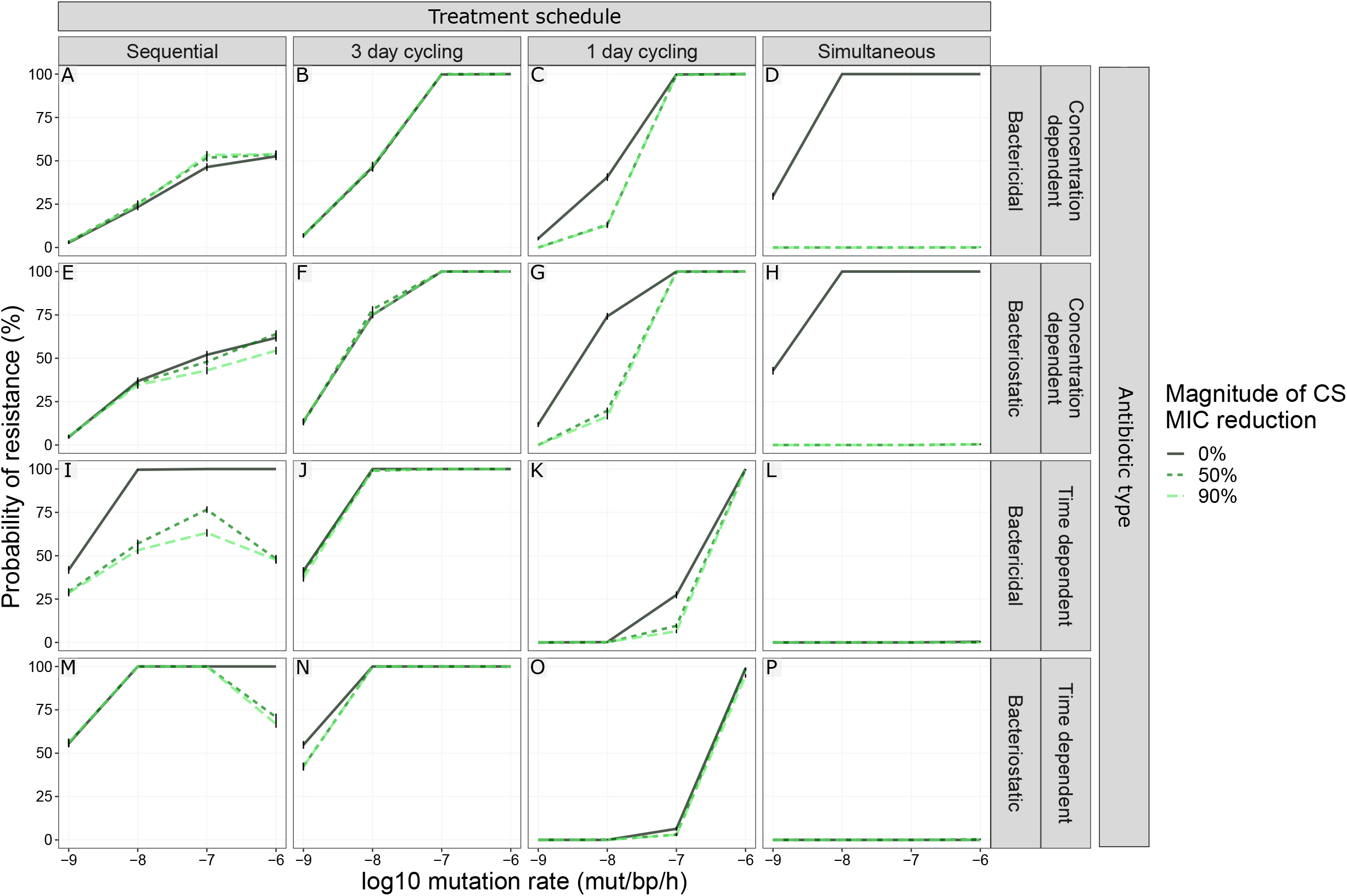
The effect of increased mutation rate for different CS magnitudes on the probability of resistance (PoR). The combined impact of mutation rate and the CS on PoR was dependent on treatment schedule. PoR of R_Any_, defined as the presence of any resistant subpopulation, was estimated at the end of treatment for treatments using different designs (columns) and antibiotic types (rows) for different mutation rates (x-axis). Each simulated scenario was realized n=500 times. Colour and line-type indicate the magnitude of reciprocal CS simulated. Data are presented as mean PoR with the error bars represent the standard error of the estimation.

## Discussion

Our theoretical analysis shows that CS can be exploited to design treatment schedules that suppress within-host development of antibiotic resistance, with CS-based treatments holding the most potential for antibiotics with narrow-therapeutic windows. Our simulations indicated that several previously unrecognised factors need to be considered to ensure optimal design of CS-based dosing regimens, which include antibiotic PD characteristics, the magnitude and reciprocity of CS effects, and the effect of fitness cost of antibiotic resistance mutations. In addition, we found that antibiotic sequence has strong impact on the success of CS-based cycling treatments. An overview of the main insights and derived design principles we obtained can be found in Supplementary Table 2.

CS-based dosing schedules have mainly considered reciprocal CS scenarios, where resistance against one antibiotic leads to increased sensitivity to a second antibiotic and vice versa^12,16^. We show, however, that one directional CS can be sufficient to supress resistance. For a one-day cycling regimen, the one-directional CS effects were nearly identical to the scenario that considered reciprocal CS (Fig. 5A vs 5B), but only when bacteria showed CS to the second drug administrated. When CS was only present for the first antibiotic (D_A_) (Fig. 5D), initial bacterial growth was extensive, thus leading to increased risk of the double resistant subpopulation emerging. Because one-directional CS relationships are much more common than reciprocal CS relationships ^9–16^, this significantly expands the number of clinical scenarios for which effective CS-based treatments can be designed.

We find that CS-based treatments show the greatest promise for antibiotics with a narrow therapeutic window. The therapeutic window of an antibiotic is defined by the drug exposure, or concentration range, leading to sufficient efficacy without associated toxicity. In the majority of our simulations, we have studied dosing schedules leading to an antibiotic steady state concentrations (C_ss_) of 1.5 x MIC_WT_ (or 0.75 x MIC_WT_ for simultaneous dosing regimens), which led to complete killing of the sensitive population but did allow emergence of resistance to occur. This concentration can be considered to reflect a narrow-therapeutic window antibiotic, *e.g*., where the antibiotic concentration required for bacterial killing is closer to the MIC because of occurrence of (severe) toxicities at higher concentrations. Indeed, for concentrations (much) higher than the MIC, or simultaneously administrating two drugs above the MIC, the benefit of CS rapidly disappears (Fig. 7). This means that especially for antibiotics with a narrow therapeutic window such as polymyxins or aminoglycosides, exploiting CS-based dosing schedules offers significant opportunities for successful antibiotic treatment while minimizing both the risks of antibiotic-related toxicity and *de novo* antibiotic-resistance development. Additionally, for simultaneously administrated antibiotics, the presence of CS could provide the possibility to lower the dosage of the individual antibiotics without decreasing efficacy.

Cycling based dosing regimens are frequently discussed as a strategy to improve antibiotic treatment when CS occurs. In our simulations, we show that for one-day cycling treatments antibiotic type (Fig. 6), directionality of CS (Fig. 5), and the identity of any pre-existing resistance subpopulation (Fig. 9) should be considered when choosing which drug to administer first. We, specifically, show that the type of the first administrated antibiotic had a larger impact on the PoR compared to the type of the second administrated antibiotic. The presences of CS to the second administrated antibiotic had a greater effect PoR compared to CS to the first administrated antibiotic. In the case of pre-existing resistance, the PoR was smaller if there was pre-existing resistance to the second administrated drug compared to the first antibiotic. These findings are consistent with previous studies showing that the probability of resistance is influenced by the sequence of antibiotics^40^, and optimized cycling sequences outperformed random drug cycling regimens^14^. Additionally, we show that one-day cycling outperforms a three-day cycling interval, both in the presence and absence of CS. This is in agreement with previous *in vitro* studies showing an advantage of shorter cycling intervals^41^. Furthermore, in the context of cycling, or alternating antibiotic treatments, consideration of the pharmacokinetics, *e.g*., the time-varying antibiotic concentrations, was found to be important because the remaining concentration of the first antibiotic administered added to the total drug effect (illustrated in Fig. 3). Therefore, the antibiotic switch contributes to a higher total drug effect than after repeated administration of the same drug, even in the absence of collateral effects. In our simulations, the impact of this increased effect is dependent on the type of the antibiotic and was shown to be especially important for time-dependent antibiotics. This highlights the importance of considering both PK and PD when designing effective antibiotic treatments, something that is overlooked when drawing conclusion regarding treatments solely based on static *in vitro* models.

To better characterize the population dynamics of pathogens in response to antibiotic treatment under presence of CS, we studied the effect of fitness costs of antibiotic resistance and mutation rates leading to antibiotic resistance. We find that introducing fitness costs had a negligible effect on PoR for the majority of the simulated CS-based treatments, with the exception of the three-day cycling using time dependent antibiotics (Fig. 8J and 8N), where introducing fitness costs improved the CS-based treatments by preventing resistant bacteria from reaching high densities before the first antibiotic switch. Typically, the PoR increased with mutation rate, which is in line with previous findings of mutator strains being associated with higher level of resistance^42,43^. For pathogens with a low mutation rate and/or administration of non-mutagenesis-inducing antibiotics (10^−9^ mut/bp/h), one-day cycling regimens and simultaneous antibiotic treatments are most relevant to benefit from CS, whereas for high mutation rates (*e.g*., 10^−6^ mut/bp/h), sequential and simultaneous antibiotic treatments are the most beneficial (Fig. 10). This means that in situations when the occurrence of mutator strains is likely (*e.g*., such as in cystic fibrosis lung infections^44^) and/or when the administered antibiotics induce mutagenesis (*e.g*., fluoroquinolones^45^), this should be considered in the design of dosing schedules. With respect to the competition between different bacterial subpopulations occurring *in vivo*, we included a bacterial carrying capacity which introduces clonal competition. During clonal competition, competition between subpopulations can lead to their suppression, *e.g*., high densities for one subpopulation can suppress the growth of a second subpopulation, even if the second population might be more fit. Treatments giving rise to clonal competition-based containment, where the selection pressure favors specific subpopulations which will in turn suppress others due to the capacity limitation of the system, have been suggested as a potential strategy to suppress AMR^46^. In our simulation, we observe a clear impact of clonal competition. When CS is present, single resistant subpopulations are unable to reach high enough levels to suppress the growth of the double resistant mutant, which allows the double mutant to take over, for some treatments. This support the value of characterizing CS-based treatments beyond the quantified summary metric of collateral effect.

Our study advances the work by Udekwu and Weiss ^22^ by explicitly comparing treatment outcomes to a base scenario without CS to determine the specific contribution of CS effects, and by performing a more systematic analysis of key drug- and pathogen specific factors that could influence optimal CS-based treatment scenarios. Additionally, we incorporated mutations as random events to capture the stochastic nature of resistance evolution, which is overlooked when using purely deterministic models. Our mathematical model was designed to facilitate identification of the primary factors driving the success or failure of antibiotic treatments in a general setting, and not for specific antibiotics and/or pathogens. We thereby did not consider factors that could further contribute to treatment outcomes for specific pathogens or antibiotics. We did not consider that more complex evolutionary mutational trajectories can occur with associated complex patters of changes in antibiotic sensitivity and MIC^47^, which are not easily definable to apply to antibiotic treatment in general. Other factors not considered include local antibiotic tissue concentrations ^48,49^, pharmacokinetic drug-drug interactions or the contribution of the immune system. We expect that such factors will not affect the specific subpopulations studied in different ways and therefore will not have a great impact on the general findings derived in this analysis.

In this analysis we assumed independent additive drug effects, thus excluding the possibility of pharmacodynamic drug-drug interactions between antibiotics, e.g., synergy or antagonism^50^. Combined drug effects can furthermore be modelled according to different null interaction assumption, including: (i) dependence of drug effects through a shared mechanism of action (Loewe additivity)^50,51^ [ref], (ii) independent drug effects with a shared maximum drug effect (Bliss independence)^52^, or (iii) fully independent additive drug effects^50^ as implemented in this paper. The choice of null interaction model, or the presence of drug interactions (synergy, antagonism) may influence treatment outcomes in particular for simultaneous treatment schedules. Although an analysis of the effect of various possible drug interactions was beyond the scope of this analysis, we do expect this will be an important factor to consider when designing CS-based treatment for specific antibiotic combinations, where specific pharmacodynamic drug interactions can be explicitly incorporated.

The developed modelling framework is applicable for design of clinical treatment designs for specific antibiotic agents and pathogens, where the model can be further expanded with additional pathogen-, drug-, and patient-specific characteristics^53^, derived from separate experimental studies and by utilizing published clinical population PK models for specific antibiotics^54,55^, which include inter-individual variability or target site concentrations at the site of infection. This would thus allow us to derive tailored CS-based dosing regimens for specific antibiotics and pathogens. Additionally, we did not evaluate how the presence of collateral resistance (CR) could impact treatment efficacy. Although such scenarios are beyond the scope of the current study, the flexibility of our developed framework allows for the incorporation of CR, and could thus serve as a tool to investigate how CR impacts treatment efficacy. Furthermore, cellular hysteresis, where non-genetic CS-like responses have been observed, may be another direction for which our modelling framework could be extended^41^.

In this study we showcase how a mathematical modelling can address questions that are difficult to answer using an experimental approach. We conclude that CS-based treatments are likely to be able to contribute in the suppression of resistance. However, the success of such treatment strategies will be dependent on careful design of a dosing schedule, and requires explicit consideration of pathogen- and drug-specific characteristics. Our developed modelling framework delineates key factors for the overall design of effective CS-informed treatments and can be used to facilitating help the design of treatments tailored to specific pathogens and antibiotic combinations. Although well-conserved CS effects remain a key requirement, we found that reciprocal CS may not be a requirement to design such dosing schedules, expanding the applicability of CS-based treatments. Such CS-based treatments appear to be most relevant for antibiotics with a narrow therapeutic window, which are also the antibiotics where within-host emergence of resistance is most likely to occur.

## Acknowledgements

We wish to acknowledge Dr. Hadi Taghvafard for helpful mathematical input, Laura Zwep for valuable discussions, and Dr. Tingjie Guo for reviewing the model code. This work was funded by ZonMW Off Road (Project number 451001033) and NWA Idea Generator (Project number NWA.1228.192.140). AL and DER were supported through the JPI-EC-AMR (Project 547001002).

## Author contributions

L.B.S.A. and J.G.C.H. designed the study; L.B.S.A. performed the data analysis; L.B.S.A., J.G.C.H., A.L., D.R. supported interpretation of results; L.B.S.A., J.G.C.H., D.R., A.L., P.H.G. wrote the paper; J.G.C.H. conceived the project; All authors reviewed the paper

## Code Availability

The model and associated code are available at https://github.com/vanhasseltlab/CS-PKPD^31^ and in Supplementary Software 1.

## Data Availability

The data simulated in this study can be generated using the available scripts. The simulated data can also be provided by the corresponding authors upon request without restrictions.

