## Supplemental materials for "Design principles of collateral sensitivity-based dosing strategies"

### Supplementary Information

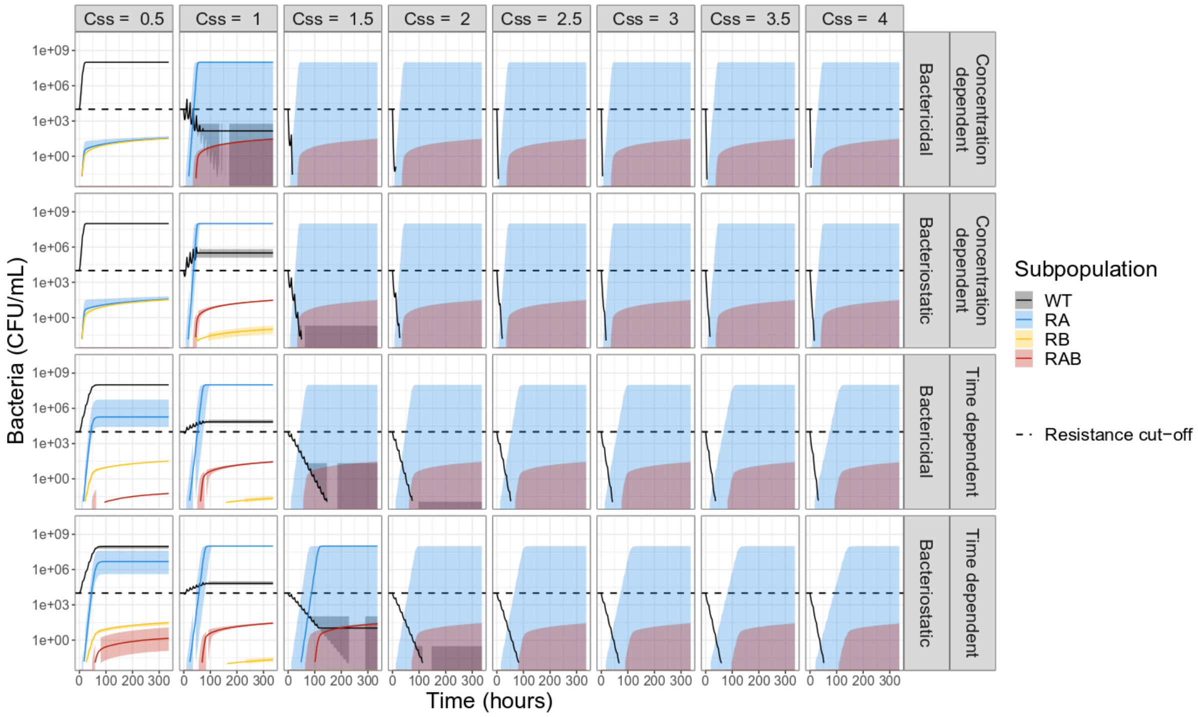


Supplementary Figure 1. **Bacterial dynamics for simulated monotherapy using different antibiotic types (rows) and steady state concentrations (Css) relating to the MIC of the wild type (WT) (columns).** These simulations shows that monotherapy required $C_{ss}$ equal to 1.5 x $MIC_{WT}$ to achieve killing of the WT, regardless of the drug type used. Each simulated scenario was realized n=500 times. Subpopulation-specific bacterial density are indicated by different colures, where the solid lines indicate the median and the shaded area covers the $5^{th}$-$95^{th}$ percentiles of the predictions. The resistance cut-off (dashed line) is used for end of treatment evaluation of resistance.

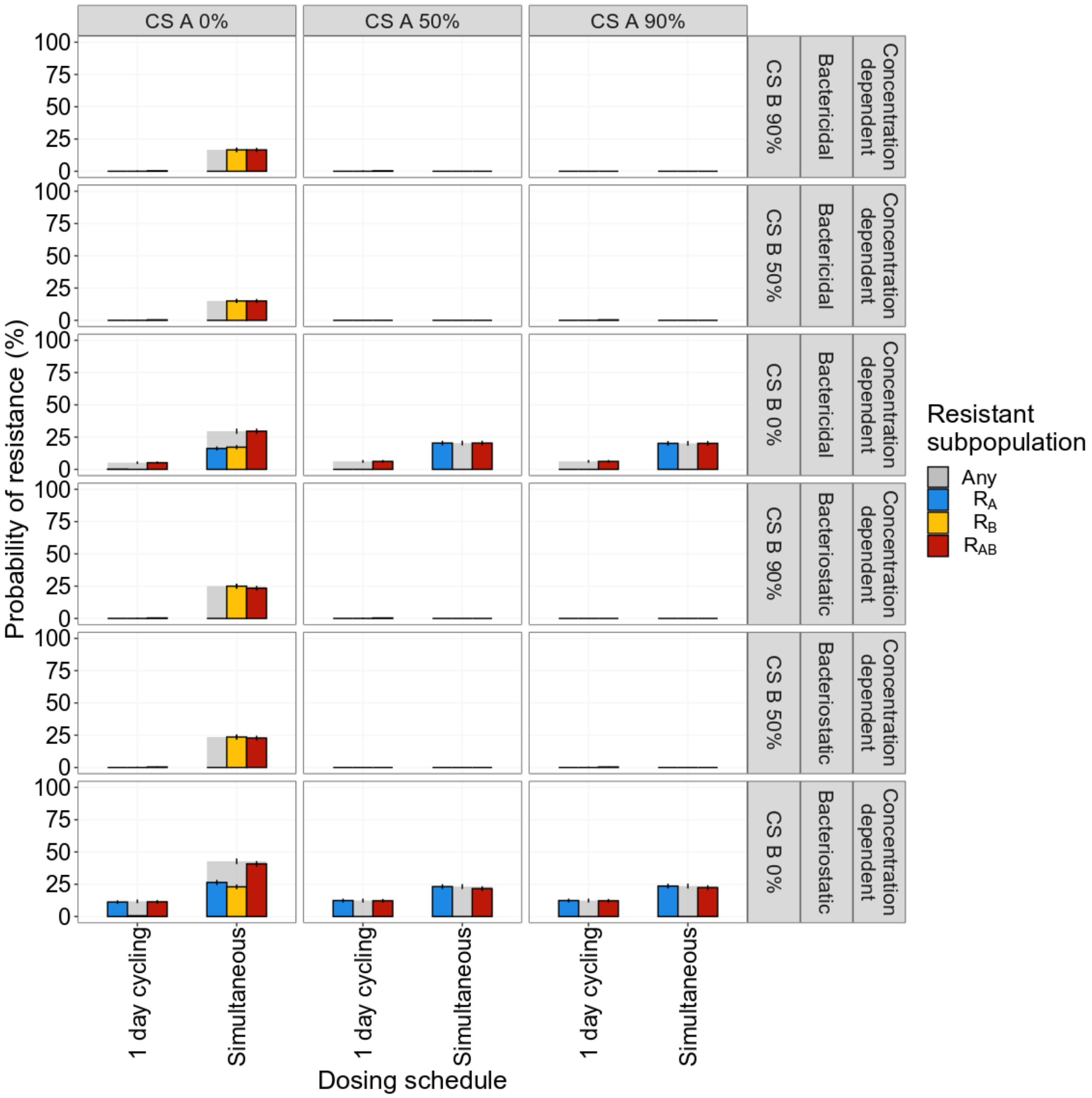


Supplementary Figure 2. **The effect of the direction or reciprocally of collateral sensitivity (CS) on end of treatment probability of resistance (PoR).** PoR was estimated at end of treatment for different CS scenarios using concentration dependent bacteriostatic or bactericidal drugs. Each simulated scenario was realized n=500 times. Subpopulation-specific PoR is indicated by different colour and $R_{Any}$ resistance, defined as the presence of any resistant subpopulation, is indicated in grey. Data are presented as mean PoR with the error bars represent the standard error of the estimation. For the one-day cycling regimen it became evident that the CS towards the second administrated drug ($AB_B$) was driving the effect, CS-based dosing using simulations administration of concentration dependent antibiotics showed that reciprocity is necessary to supress overall resistance.

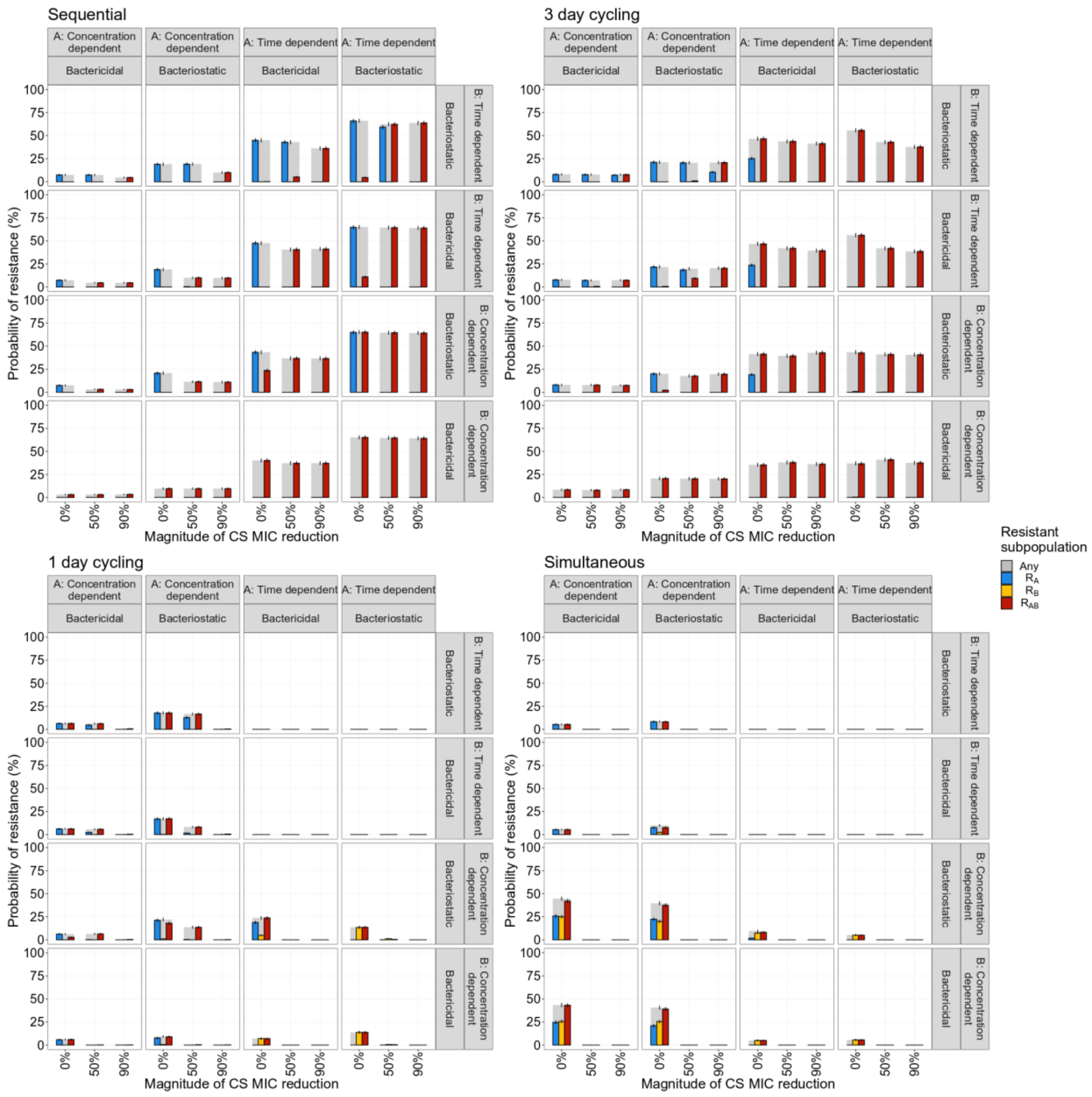


Supplementary Figure 3. **The effect of using different antibiotic combinations during treatments in relation to different levels of collateral sensitivity (CS) on the probability of resistance (PoR) at the end of treatment.** PoR was estimated at the end of treatment for different treatment schedules with different antibiotic combinations. Each simulated scenario was realized n=500 times. Subpopulation-specific PoR is indicated by different colour and $R_{Any}$, defined as the presence of any resistant subpopulation, is indicated in grey. Data are presented as mean PoR with the error bars represent the standard error of the estimation.

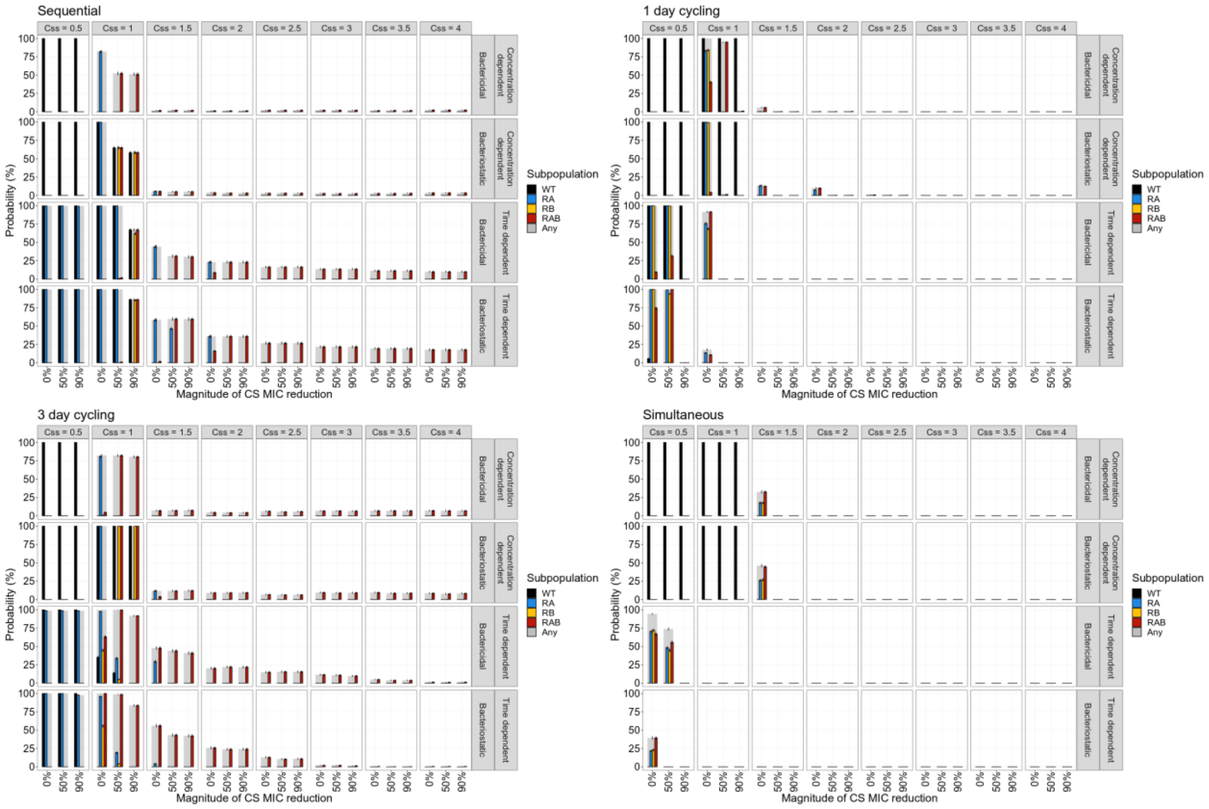


Supplementary Figure 4. **The effect of antibiotic steady state concentrations ($C_{ss}$) in relation to different levels of collateral sensitivity (CS) on the probability of resistance at the end of treatment (PoR).** $C_{ss}$ was expressed as factor difference from the $MIC_{WT}$. PoR of $R_{Any}$, defined as the presence of any resistant subpopulation, was estimated at the end of treatment for treatments using different designs and antibiotic types (rows). Each simulated scenario was realized n=500 times. Subpopulation-specific PoR is indicated by different colour and $R_{Any}$ is indicated in grey. Data are presented as mean PoR with the error bars represent the standard error of the estimation.

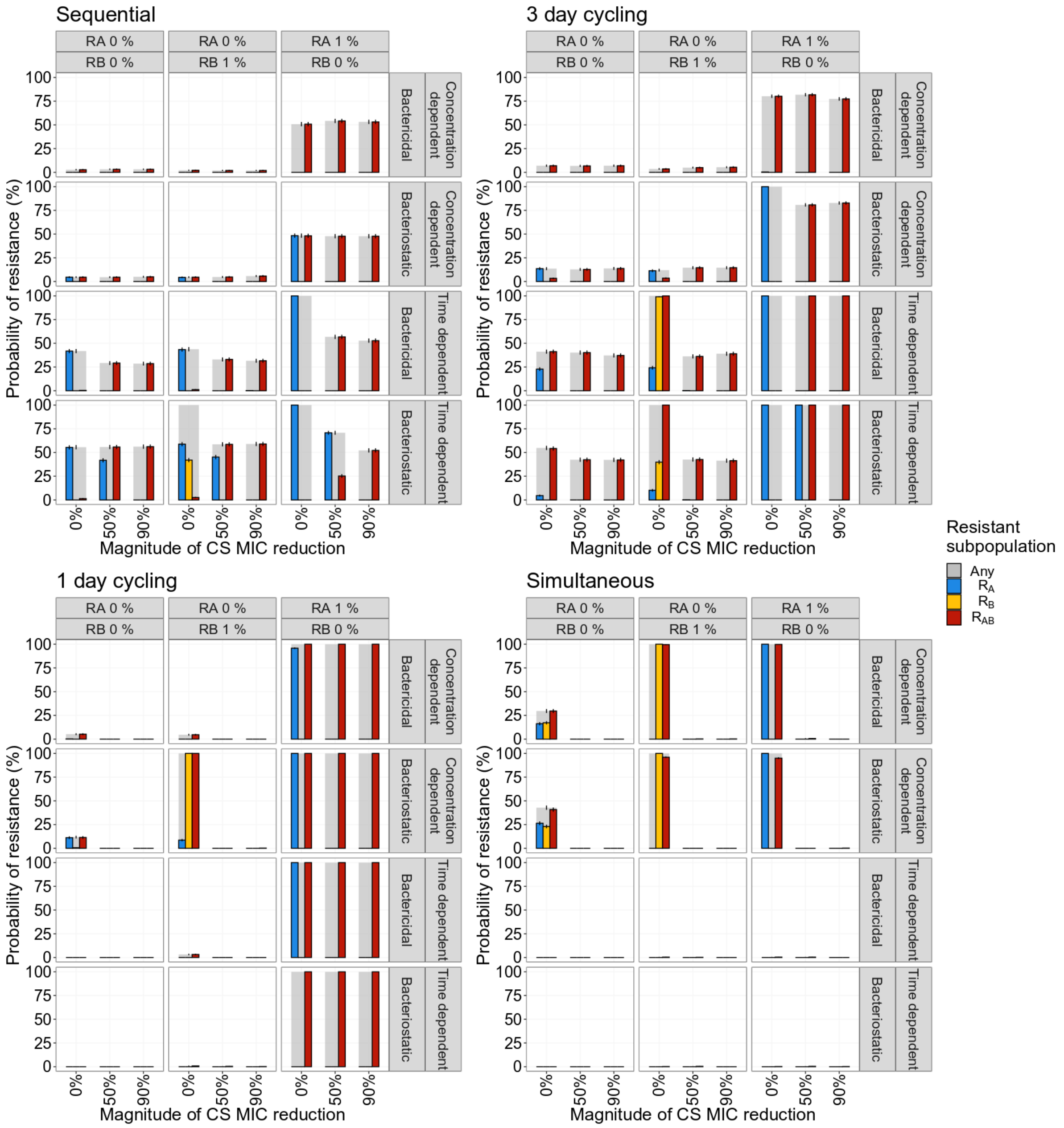


Supplementary Figure 5. **The effect of pre-existing resistant mutants for different magnitudes of collateral sensitivity on the probability of resistance (PoR).** PoR was estimated at the end of treatment for different scenarios of low levels of pre-existing resistance (columns) and antibiotic types (rows). Each simulated scenario was realized n=500 times. Subpopulation-specific probability of resistance is indicated by colour and PoR of $R_{Any}$, defined as the presence of any resistant subpopulation, is indicated in grey. Data are presented as mean PoR with the error bars represent the standard error of the estimation.

Supplementary Table 1. **Simulation scenarios evaluated and associated pathogen- and pharmacological factors studied**

| **Scenario** | **Pathogen factors** | | | | **Treatment factors** | |
|---|---|---|---|---|---|---|
| | Collateral sensitivity (%) | Fitness cost per mutation (%) | Mutation rate (mut/bp/h) | Pre-existing resistance | PD parameters | Steady state concentration* |
| **1: Treatment design** | Symmetric reciprocal: 50 or 90 | No | $10^{-9}$ | No | Same-type combinations: $G_{\min,D_A} = G_{\min,D_B}$ and $\mathrm{Hill}_{D_A} = \mathrm{Hill}_{D_B}$ | 1.5x $MIC_{WT}$ |
| **2: Directionality of CS** | One directional or asymmetric reciprocal: 50 or 90 | No | $10^{-9}$ | No | Same-type combinations: $G_{\min,D_A} = G_{\min,D_B}$ and $\mathrm{Hill}_{D_A} = \mathrm{Hill}_{D_B}$ | 1.5x $MIC_{WT}$ |
| **3: Drug type combinations** | Symmetric reciprocal: 50 or 90 | No | $10^{-9}$ | No | Different types combined: $G_{\min,D_A} \neq G_{\min,D_B}$ and/or $\mathrm{Hill}_{D_A} \neq \mathrm{Hill}_{D_B}$ | 1.5x $MIC_{WT}$ |
| **4: Therapeutic window** | Symmetric reciprocal: 50 or 90 | No | $10^{-9}$ | No | Same-type combinations: $G_{\min,D_A} = G_{\min,D_B}$ and $\mathrm{Hill}_{D_A} = \mathrm{Hill}_{D_B}$ | 0.5-5x $MIC_{WT}$ |
| **5: Fitness cost** | Symmetric reciprocal: 50 or 90 | Yes: 10, 20, 30, 40, or 50 | $10^{-9}$ | No | Same-type combinations: $G_{\min,D_A} = G_{\min,D_B}$ and $\mathrm{Hill}_{D_A} = \mathrm{Hill}_{D_B}$ | 1.5x $MIC_{WT}$ |
| **6: Pre-existing resistance** | Symmetric reciprocal: 50 or 90 | No | $10^{-9}$ | Yes: 1% $R_A$ or 1% $R_B$ | Same-type combinations: $G_{\min,D_A} = G_{\min,D_B}$ and $\mathrm{Hill}_{D_A} = \mathrm{Hill}_{D_B}$ | 1.5x $MIC_{WT}$ |
| **7: Mutation rate** | Symmetric reciprocal: 50 or 90 | No | $10^{-9}$, $10^{-8}$, $10^{-7}$, or $10^{-6}$ | No | Same-type combinations: $G_{\min,D_A} = G_{\min,D_B}$ and $\mathrm{Hill}_{D_A} = \mathrm{Hill}_{D_B}$ | 1.5x $MIC_{WT}$ |

CS = Collateral sensitivity; PD = Pharmacodynamics; $G_{\min,Di}$ = type of antibiotic effect; $\mathrm{Hill}_{Di}$ = driver of antibiotic effect; $MIC_{WT}$ = 1 mg/L

*divided by 2 for simultaneous dosing regimens

Supplementary Table 2. **Overview of derived design principles based on evaluated simulation scenarios.**

| Scenario | Design principles | Figure |
|---|---|---|
| **1a: Treatment design - Drug type** | • The choice of drug type, *i.e.,* bactericidal or bacteriostatic, and time- or concentration-dependent, influences the probability of resistance.<br>• The driver of the effect (time- vs concentration-dependent) has a larger impact than the type of effect (bactericidal vs bacteriostatic). | 4 |
| **1b: Treatment design - Treatment schedule** | • The choice of treatment schedule, *i.e.,* sequential, cyclic or simultaneous administration, influence the probability of resistance. | |
| **2: Directionality of CS** | • One-directional CS can be enough to supress antibiotic resistance.<br>• Antibiotic cycling is more effective than simultaneous administration for one-directional CS.<br>• Therapy should be initiated with the antibiotic not showing CS. | 5, S2 |
| **3: Drug type combinations** | • Therapy should be initiated with a time-dependent antibiotic if both a concentration- and a time-dependent antibiotic are used. | 6, S3 |
| **4: Therapeutic window** | • CS-based treatments show the most potential for antibiotics with a narrow therapeutic window. | 7, S4 |
| **5: Fitness cost** | • Three-day cycling treatments using time-dependent antibiotics benefit from the fitness cost of antibiotic resistance mutations. | 8 |
| **6: Pre-existing resistance** | • Cycling therapy should be initiated with the antibiotic for which no low-level pre-existing resistance is present. | 9, S5 |
| **7: Mutation rate** | • Low mutation rate: use one-day cycling or simultaneous antibiotic administration.<br>• High mutation rate: use sequential or simultaneous antibiotic administration. | 10 |

CS=Collateral sensitivity; S = Supplementary Figure
